# Human herpesvirus 8 molecular mimicry of ephrin ligands facilitates cell entry and triggers EphA2 signaling

**DOI:** 10.1101/2021.03.26.437132

**Authors:** Taylor P. Light, Delphine Brun, Pablo Guardado-Calvo, Riccardo Pederzoli, Ahmed Haouz, Frank Neipel, Felix Rey, Kalina Hristova, Marija Backovic

## Abstract

Human herpesvirus 8 (HHV-8) is an oncogenic virus that enters cells by fusion of the viral and endosomal cellular membranes in a process mediated by viral surface glycoproteins. One of the cellular receptors hijacked by HHV-8 to gain access to cells is the EphA2 tyrosine kinase receptor, and the mechanistic basis of EphA2-mediated viral entry remains unclear. Using X-ray structure analysis, targeted mutagenesis and binding studies, we here show that the HHV-8 envelope glycoprotein complex gH/gL binds with sub-nanomolar affinity to EphA2 via molecular mimicry of the receptor’s cellular ligands, ephrins, revealing a pivotal role for the conserved gH residue E52 and the amino-terminal peptide of gL. Using FSI-FRET and cell contraction assays, we further demonstrate that the gH/gL complex also functionally mimics ephrin ligand by inducing EphA2 receptor association via its dimerization interface, thus triggering receptor signaling for cytoskeleton remodeling. These results now provide novel insight into the entry mechanism of HHV-8, opening avenues for the search of therapeutic agents that could interfere with HHV-8 related diseases.

## INTRODUCTION

Human herpesvirus 8 (HHV-8), also known as Kaposi’s sarcoma-associated virus, is a member of *Rhadinovirus* genus that belongs to the *Gammaherpesvirinae* subfamily of *Herpesviridae* ^1^. HHV-8 is an oncogenic virus and etiological agent of Kaposi’s sarcoma (KS), malignancy of endothelial cells named after the Hungarian dermatologist who first described the disease in 1872 ^2^. Because KS has different clinical manifestations, two main forms are distinguished - the classic KS that is a relatively indolent and rare tumor, appearing as skin lesions mostly in elderly men, and the epidemic or HIV-associated KS, an aggressive form that spreads extensively through skin, lymph nodes, intestines and lungs ^3^. The KS affects up to 30% of untreated HIV-positive individuals ^4,5^, and is nowadays one of the most frequent malignancies in men and children in subequatorial African countries ^6^.

Behind the ability of HHV-8 to spread to diverse tissues lies its wide tropism demonstrated *in vivo* for epithelial and endothelial cells, fibroblasts, B- and T-lymphocytes, monocytes, macrophages and dendritic cells (reviewed in ^7^). As other herpesviruses, HHV-8 attaches to cells via its envelope glycoproteins that engage in numerous low affinity interactions with ubiquitous cellular factors such as heparan sulfate proteoglycans ^8^. Following these initial contacts, the viral envelope glycoprotein(s) bind to specific cell receptor(s) in an interaction that provides trigger for membrane fusion. The envelope glycoproteins B (gB) and the non-covalent heterodimer made of glycoproteins H and L (gH/gL) constitute the conserved core fusion machinery of all herpesviruses, whose function is to induce merger of the viral and cellular membranes. The release of herpesvirus capsids occurs at the level of plasma membrane or as in the case of HHV-8, after endocytosis, by fusion of the viral and endosomal membranes ^9^. In all herpesviruses gB is the fusogen protein, while gH/gL plays a role in regulation of gB activity ^10^. What is particular to HHV-8 is the simultaneous employment of several viral glycoproteins - HHV-8-specific K8.1A glycoprotein, as well as the core fusion machinery components – that engage diverse cellular receptors (gB binds to integrins and DC-SIGN, gH/gL to EphA receptors) increasing the HHV-8 target repertoire and providing the virus with a set of tools for well-orchestrated entry (reviewed in ^11^).

The X-ray structures of gH/gL from alpha-, beta- and gammaherpesviruses ^12–15^, revealed a tightly bound protein complex, in which the gH ectodomain, composed of four domains forms a tight complex with gL via its N-terminal, membrane-distal domain. Despite low gH and gL sequence conservation across the herpesvirus family, the structures are remarkably similar, indicating conservation of a function. The current fusion model posits that upon binding of the virus to the cellular receptor (via gH/gL, gB, or a virus-specific glycoprotein), gH/gL in a still unknown way relays the signal and switches gB fusion activity on, setting membrane fusion in motion ^16^. As mentioned above, HHV-8 gH/gL also functions as a receptor binding protein that interacts with cellular Eph receptors that belong to a superfamily of transmembrane tyrosine kinases, named Eph for their expression in <u>e</u>rythropoietin-<u>p</u>roducing human <u>h</u>epatocellular carcinoma cell line ^17^.

The physiological ligands of Eph receptors are membrane-tethered proteins called ephrins (acronym for <u>Eph</u> family <u>r</u>eceptor <u>in</u>teracting proteins). Ephrin-A ligands are attached to the membrane by glycosylphosphatidylinositol anchors and bind to Eph receptors classified as type-A Eph receptors, while ephrin-B ligands possess a transmembrane and an intracellular PDZ domain and interact with receptors designated as Eph type-B receptors ^18,19^. There are 5 ephrin-A ligands and 9 Eph type-A receptors in humans. HHV-8 binds with the highest affinity to EphA2, and less to the related EphA4 and EphA7 receptors ^20–22^. In addition, the EphA2 receptor serves as a receptor for two other gammaherpesviruses - human herpesvirus 4 (HHV-4 also known as Epstein-Barr virus (EBV) and rhesus monkey rhadinovirus ^23,24^. The interactions are in all cases established via gH/gL. Unrelated pathogens such as Hepatitis C virus, some bacteria and yeast also depend on EphA2 receptor for entry ^25–28^.

Eph receptor-ephrin ligand interactions mediate short-distance cell-cell communications and lead to cytoskeleton rearrangements and rapid changes in cell mobility and / or morphology ^29^. Some of the typical outcomes of ephrin-A1 ligand activation of EphA2 receptor are cell retraction ^30–33^ and endocytosis of receptor-ligand complexes ^34^. These processes are especially active and important during development, and in adulthood many of the same circuits get repurposed for functions in bone homeostasis, angiogenesis, synaptic plasticity (reviewed in ^29^). Since motility and angiogenesis contribute to tumorigenesis and other pathologies, Eph receptors and ephrin ligands are in the spotlight as targets for therapeutic intervention ^35,36^.

EphA2 was identified as HHV-8 entry receptor by Hahn *et al.* who showed that deletion of the EphA2 gene abolished infection of endothelial cells, and that binding of gH/gL to EphA2 on cells led to increased EphA2 phosphorylation and endocytosis facilitating viral entry. The presence of the intracellular kinase domain was shown to be important for HHV-8 entry in epithelial 293 cells ^20^. In this respect, HHV-8 gH/gL does not play the role of a classical herpesvirus receptor binding protein, directly activating gB upon binding to a cellular receptor to induce fusion of the viral and plasma membranes, such as gD in alphaherpesviruses. The merger of the HHV-8 and cellular membranes occurs at low pH within an endosome, and is spatially and temporally separated from the gH/gL interactions with EphA2 that occur at the cell surface. HHV-8 gH/gL instead activates EphA2 receptors that initiate signaling pathways leading to rapid internalization of the virus and cytoskeletal rearrangements that create a cellular environment conducive for the virus and capsid intracellular transport ^37^.

All Eph receptors contain an elongated ectodomain made of - as beads on a string - a ligand-binding domain (LBD), a cysteine-rich domain (CRD) and two fibronectin (FN) domains, followed by a transmembrane anchor, a short juxtamembrane region containing several conserved tyrosine residues, an intracellular Tyr kinase domain, a sterile alpha motif (SAM) that has a propensity to oligomerize, and a PDZ domain involved in protein-protein interactions ^38^. The LBD consists of a rigid, conserved β-sandwich, decorated with variable loops that dictate ligand specificity ^39,40^. In a similar fashion, the protruding flexible loops of ephrin ligands are arranged around a compact, eight-stranded central β-barrel ^41^. We employ the accepted nomenclature for the secondary structure elements for Eph receptors ^42^ and ephrin ligands throughout the text ^41^. In both cases single letters designate β-strands and helices, and double letters are used to label loops that connect the secondary structure elements (Fig. S1). The ephrin ligand interactions with the Eph receptors are driven by an 18-residue long and mostly hydrophobic loop that connects strands G and H - the GH loop - of the ligand, which inserts into a complementary hydrophobic cavity presented at the surface of the receptor LBD ^42^. The ephrin ligand GH loop carries a conserved E119 that establishes polar interactions, critical for high affinity binding, with a strictly conserved R103 on the loop of the Eph type-A receptors. This R103 happens to be in the loop connecting strands G and H in EphA receptors and is also designated as GH loop ^43^. To avoid confusion, we use superscripts to indicate the molecule of the residue or feature described (R103^EphA2^, E119^ephrin^, GH^EphA2^, GH^ephrin^ etc).

At the molecular level, as in the case of other receptor tyrosine kinases, ephrin ligand binding induces oligomerization of Eph receptors, promoting trans-phosphorylation and signal transduction into the cell ^19^. Structural and functional studies revealed that ephrin ligand binding to Eph receptors results first in formation of so-called tetramers made of two ‘Eph-ephrin’ complexes in which each Eph receptor interacts with 2 ephrin molecules - its cognate ligand with high affinity, and via low affinity interactions with ephrin from the other complex ^18^ (Fig. S2). In such tetrameric arrangement that is brought about by ligands, two Eph receptors are arranged into a dimer (also known as homodimer), stabilized by contacts at the EphA2 LBD dimerization interface (DIN). Eph-ephrin tetramers can polymerize into clusters via the interface in the downstream CRD referred to as the clustering interface (CIN) ^44–46^. Since the clustering involves an EphA2 region outside of the LBD, this interface is known as ligand independent. Indeed EphA2 receptor was shown to exist on cells in the absence of ligand in a monomer-dimer equilibrium ^47^, with the unliganded dimers stabilized via the CIN (Fig. S2).

The cellular response to EphA2 receptor activation is ligand- and cell-type dependent and modulated by factors such as the spatial distribution of the receptor in the membrane ^48^, residues in the intracellular domain that are phosphorylated, size and type of the EphA2 receptor oligomers, to just name some ^29^. EphA2 receptor dimers were shown to be already active ^49^, and there is evidence that oligomers made of 6-8 EphA2 receptor molecules activate, while larger aggregates may dampen the signaling in some cell types ^50^. Different ligands (monomeric, dimeric ephrin-A1, and agonist or antagonist peptides) were shown to stabilize distinct dimeric or oligomeric EphA2 receptor assemblies (Fig. S2), further indicating that the signaling properties may be defined by the nature of the EphA2 dimers and oligomers ^47,51^.

The structures of several Eph receptor – ephrin ligand complexes have been determined ^39,40,42,44,45^, but how viral antigens such as HHV-8 gH/gL interact with EphA2 was completely unknown until recently. The 3.2Å X-ray structure of a HHV8 gH/gL-EphA2 complex was published while we were preparing this manuscript ^52^. Our goal has been to explore the events that emulate the early stages of HHV-8 entry. We south to obtain the structural details on the gH/gL-EphA2 complex, and to determine if and how HHV-8 gH/gL affects assembly of EphA2 receptors in the membranes of living cells, taking advantage of the FSI-FRET system that allows following lateral interactions of membrane proteins *in vivo* ^53^. We report here a 2.7Å resolution X-ray structure of the HHV-8 gH/gL ectodomain bound to the LBD of EphA2 together with results of structure-guided mutagenesis and cell-based studies that highlight E52^gH^ as a key residue for high-affinity binding to the receptor. Based on our analyses and the similarities we observed between the binding modes of gH/gL and the ephrin ligand to EphA2, we provide evidence that this structural similarity extends into functional mimicry - soluble gH/gL induces cell contraction and stabilizes the same type of EphA2 receptor dimers on cells, as was shown for the ephrin-A1 ligand ^47^. This is the first time that a non-ephrin protein in its monomeric form has been shown to activate Eph receptors, underscoring that signaling via EphA2 dimers, and not large oligomers, may be more common than thought. The results presented here now lay a path for further exploration of downstream events and the investigation of whether HHV-8 activation of Eph receptors may play a role beyond ensuring a productive infection, for example contributing to virus oncogenicity/oncogenic transformation of the cell.

## RESULTS

### Structure determination of the EphA2 LBD - gH/gL tertiary complex

Recombinant HHV-8 gH/gL ectodomains and EphA2 LBD were expressed in insect cells and the proteins were purified as described in detail in Material and Methods (MM). To maximize the tertiary complex (gH/gL-bound to EphA2 LBD) formation, the gH/gL was mixed with an excess of LBD which was then removed by size exclusion chromatography. Multiangle light scattering measurements ^54^ demonstrated a 1:1:1 tertiary complex stoichiometry for gL/gH bound to EphA2 LBD, as well as to the EphA2 ectodomain (Fig. S3).

The tertiary complex crystallized in an orthorhombic space group (C 2 2 2_1_) and the crystals, which contained one molecule per asymmetric unit, diffracted to 2.7Å. The initial phases were calculated by molecular replacement using the EphA2 LBD (PDB: 3HEI) and an HHV-8 gH/gL theoretical model derived from the EBV gH/gL X-ray structure (PDB: 3PHF) using the Phyre2 program for protein modelling and structure prediction ^55^. The partial molecular replacement solution was extended by iterative cycles of auto- and manual building as explained in MM. The final map displayed clear electron density for residues 27-200 of EphA2, residues 21-128 for gL, and residues 35-696 gH with the exception of several short regions of poor density that precluded unambiguous placement of the polypeptide chain (Fig. 1A). The N-terminus of gL contained two additional residues (Arg and Ser) carried over from the expression vector. Around 40 gL residues at its C-terminus were not resolved and are likely to be disordered as predicted by IUPred2A server ^56,57^. The atomic model was refined to a R_work_/R_free_ of 0.22/24 (Table S1).

**Figure 1:**
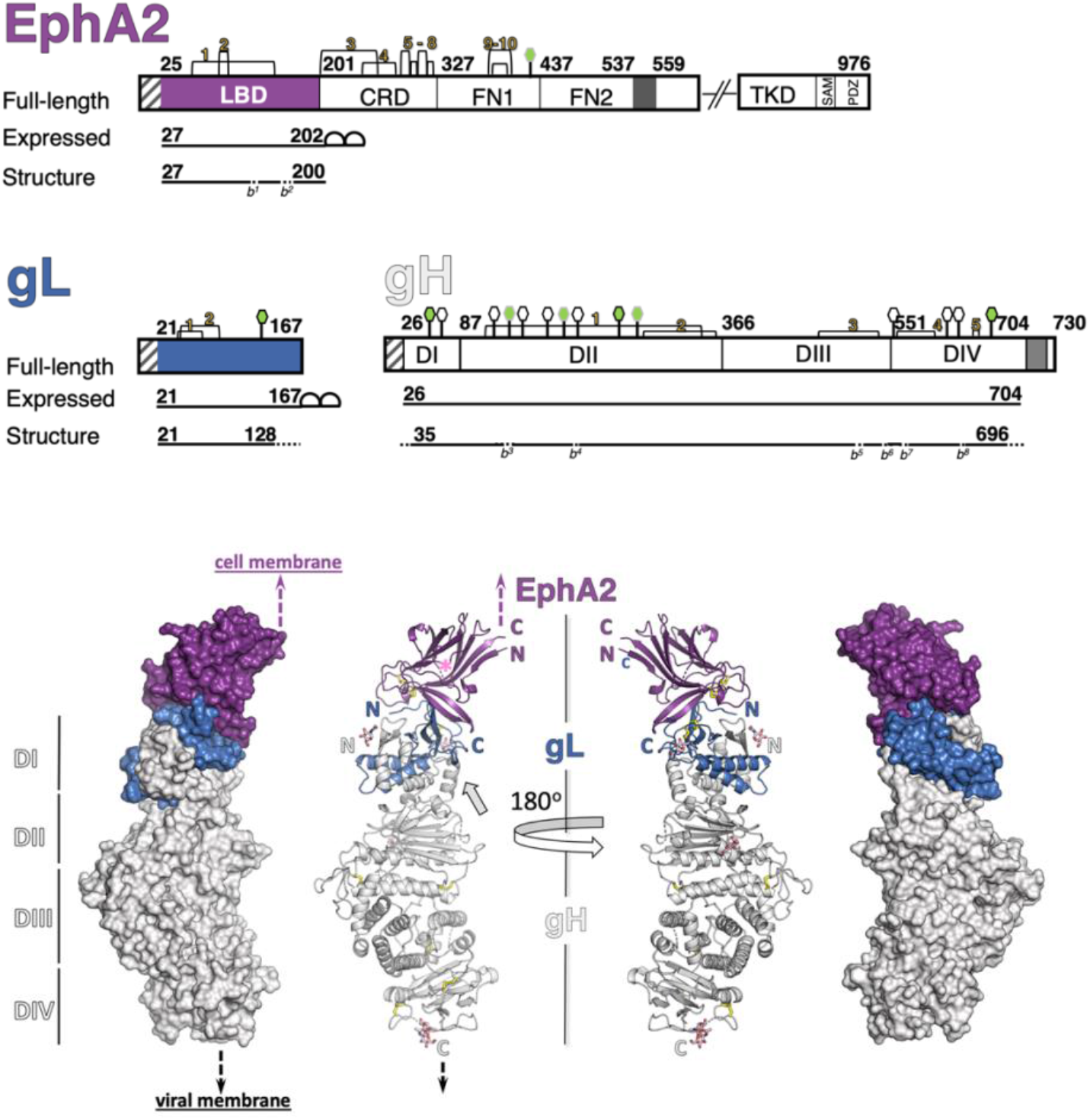
Schematic representation of HHV-8 gH, gL and EphA2 and structure of the gH/gL-EphA2 LBD complex. **A)** Schematic representation of the EphA2 receptor, HHV-8 gL and gH, highlighting the protein segments that were expressed as recombinant proteins for crystallization, and the residues resolved in the structure. The short fragments that could not be built in EphA2 LBD and gH because of the poor electron density are marked with dotted lines and labeled as breaks (*b*) corresponding to the missing residues: *b^1^* (111), *b^2^* (148-162) in EphA2 LBD, and *b^3^*(127-131), *b^4^*(212-216), *b^5^*(521-526), *b^6^*(547-550), *b^7^*(558-559), *b^8^*(627-629) in gH. The disulfide bonds are indicated with yellow numbers, and N-linked glycosylation sites with hexagons (green with black border - built in our structure; green with gray border - built in the PBD accession code 7CZF [1]; white, with black borders - remaining, predicted sites). Signal peptides at the start of each protein are represented as white boxes with grey lines, transmembrane anchor domains in EphA2 and gH as dark grey boxes, and double strep tag for affinity purification on gL and EphA2 LBD as half circles. **B)** The structure of the tertiary complex is represented as molecular surface and cartoon model (EphA2 LBD in purple, gL in blue and gH in grey). The N- and C-termini of each protein are labelled with letters “N” and “C”, respectively. The four domains of gH are marked with roman numbers on the left side, and putative locations of the viral and cellular membranes with dashed arrows (black and purple, respectively). The hinge / linker region on gH is indicated with a grey arrow, and putative position of the unresolved J helix in the LBD with a pink * symbol.

### The gH/gL-EphA2 LBD complex structure

The tertiary complex forms an extended structure 15 nm long and around 4.6 nm across its widest part, in the gH region (Fig. 1B). EphA2 LBD adopts a jelly-roll fold as originally described ^39^ – its N- and C-termini point in the same direction and away from the gH/gL binding site, consistent with the expected location of the remaining EphA2 domains. Two antiparallel 5-stranded β-sheets pack into a compact β-sandwich, with loops of different lengths connecting the strands. The HI^EphA2^ loop is well ordered and forms the DIN, while the JK^EphA2^ loop, which carries a short J’ helix, is not resolved in our tertiary complex structure likely due to its already reported structural plasticity ^40^ and / or displacement by gL (Fig.1B). Apart from the JK^EphA2^ loop, the EphA2 LBD does not change conformation upon binding to gH/gL.

The gH/gL complex has an architecture already described for other herpesvirus orthologs – the γ-herpesvirus EBV gH/gL ^15^, β-herpesvirus human CMV ^13^, and α-herpesviruses HSV-2, PrV and VZV ^14,58,59^. The N-terminal domain I (DI) of gH is separated by a linker (hinge) helix from the rest of the ectodomain (domains II, III and the membrane-proximal domain IV) (Fig. 1B). The HHV-8 gH/gL resembles the most its EBV counterpart, consistent with the highest sequence conservation between the two, followed by the β-herpesvirus CMV gH/gL complex and less so the α-herpesvirus complexes (Fig. S4). The rmsd values and Z-scores calculated from the superimposition of individual gH domains and gL are given in Fig. S4 (superimposing the entire gH/gL ectodomains is not informative because of the different orientations of the domains with respect to each other). Clear electron density was observed at 4 N-linked glycosylation sites (N46^gH^, N267^gH^, N688^gH^, and N118^gL^) allowing placement of 1 or 2 N-acetylglucosamine residues.

### Binding interface between gH/gL and EphA2

The EphA2 LBD and gH/gL form an intricate interface structure made of a seven-stranded mixed β-sheet containing strands contributed by all 3 proteins. The N-terminal segment of gH co-folds with gL forming a mixed five-stranded β-sheet composed of two gH and three gL β-strands. The third gL β-strand further engages in contacts with the D β-strand of EphA2 (Fig. 2A).

**Figure 2:**
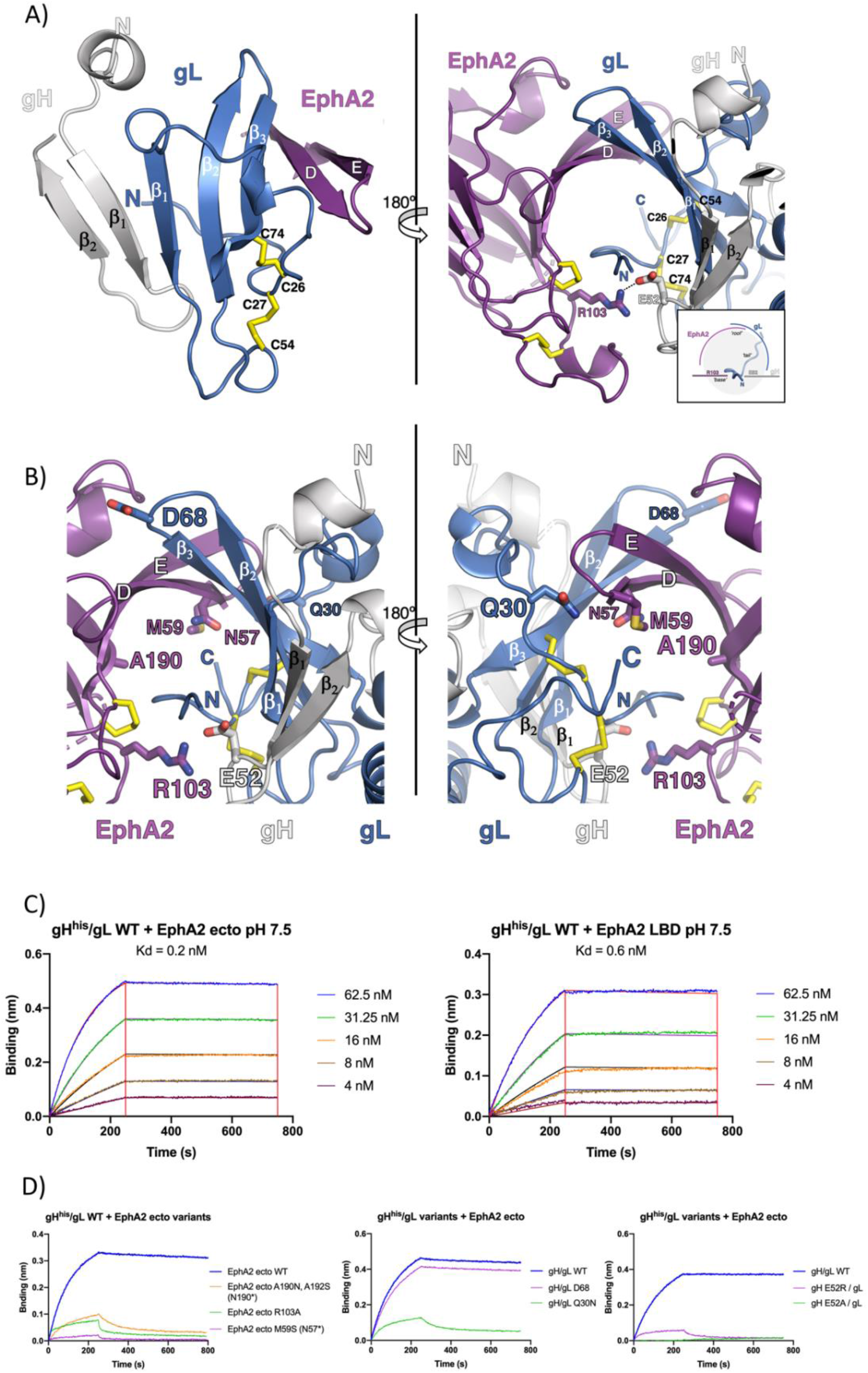
The binding interface between EphA2 LBD and HHV-8 gH/gL. **A)** The mixed β-sheet formed by strands of gH, gL and EphA2. The strands in gH and gL are labeled as β_number_, while the EphA2 LBD strands are marked using the single-letter nomenclature assigned for the first solved structure of the EphB2 LBD 1KGY [2]. The same coloring scheme as in Figure 1 is applied (left panel). View are the gH/gL and EphA2 binding interface from the other side with the inlet illustrating the channel formed by the EphA2 and gL strands (‘roof’) that accommodates the gL N-terminal ‘tail’, reinforced by polar interactions between R103^EphA2^ and E52^gH^ (‘base’) (right panel). **B)** Locations of the point mutations introduced in EphA2 (R103A) and gH (E52A, E52R), and N-linked glycosylation sites in EphA2 (N57, A190N) and gL (Q30N, D68N) are indicated, and their side chains are shown as sticks. The same coloring scheme as in Figure 1B is applied. **C)** Sensorgrams recorded for WT EphA2 ectodomain of LBD binding to immobilized gH/gL by biolayer interferometry. A series of measurements using a range of concentrations for EphA2 ectodomain and LBD, respectively, was carried out to obtain the Kd for the WT proteins. Experimental curves (colored traces) were fit using a 1:1 binding model (black traces) to derive equilibrium *K*_d_ values. **D)** Sensorgrams recorded for EphA2 variants binding to immobilized gH/gL variants by biolayer interferometry. Single experimental curves obtained for EphA2 ectodomain concentration of 62.5 nM plotted to show the effect of the, respectively, EphA2 mutations, gL mutations, and gH mutations on binding.

gL binds to the EphA2 LBD via its N-terminal segment (residues 21-30) and residues from its β2 and β3 strands (Fig. 2A) (the full list of contact residues is given in Table S2 and is represented in Fig. S5). The gL N-terminal segment is restrained by C26 and C27 that form disulfide bonds with C74 and C54, respectively. Immediately upstream this anchoring point there is an elongated, hydrophobic ‘tail’ (residues 21-25) that inserts into a hydrophobic channel formed by the EphA2 strands D and E and gL strands β2 and β3 (the ‘roof’). The buried surface area for the gL residues 19-32 is around 480 Å^2^. Of the 14 hydrogen bonds formed between gL and EphA2, 7 are contributed by the gL N-terminal segment, 6 by strand β3 and 1 by the C-terminal η4 Asn128.

Below the gL ‘tail’, the single gH residue that makes contacts with EphA2 - E52^gH^ - forms a salt bridge with R103^EphA2^, clamping the bottom of the tunnel (the ‘base’). E52^gH^ is also involved in polar interactions with residues V22^gL^, H47^gL^ and F48^gL^, thus being a center point (a hub) interlaying gL and EphA2 (Table S2).

### Biolayer Interferometry (BLI) analyses of EphA2 and gH/gL interactions in solution

To investigate the role of the gH/gL and EphA2 residues implicated in the interactions observed in the crystal structure, we tested a series of mutants that were conceived to induce large perturbations into gL and EphA2 by introducing N-glycosylation sites. We resorted to such drastic changes because most EphA2 point mutations already tested in immunoprecipitation assays had only moderate effects on gH/gL binding ^60^. The point mutation R103A^EphA2^ has been reported to abolish the binding to gH/gL ^52^ and served as a positive control. Since E52^gH^ is the only gH residue contacting EphA2, we also introduced point mutations E52A^gH^ and E52R^gH^ to specifically target this site.

The variants with the following N-glycosylation sites were generated: N57^EphA2^, N190^EphA2^, N30^gL^ or N68 ^gL^, and point mutant variants R103A^EphA2^, E52A^gH^ bound to gL (E52A^gH^/ gL) and E52R^gH^/ gL (Fig. 2B). The recombinant proteins were expressed in mammalian cells. The introduced sites N30^gL^ and N68 ^gL^ were glycosylated as clearly observed by gL shift to a higher molecular weight on SDS-PAGE gels, while the change in the migration was harder to detect for EphA2 ectodomains possibly because its larger size and small difference introduced by an additional glycosylation. The N190^EphA2^ mutation was already reported to perturb the interactions with ephrin ligands ^44^. The constructs were engineered so that gL contained a strep tag for complex purification, as before, and gH contained a histidine tag at C-terminus for immobilization onto BLI sensors via the end distal to the EphA2 binding site (Fig. S6). Further details on protein production and BLI parameters are given in MM.

We determined the dissociation constant (Kd) <1nM by doing a series of BLI measurements for wild type (WT) gH/gL binding to the EphA2 ectodomain (res. 27 - 534) or its LBD (res. 27-202) (Fig. 2C). The low Kd observed for the WT proteins was dominated by a slow k_off_ rate. We obtained a Kd in the subnanomolar range when the measurements were done at pH 5.5 (Fig. S7A), or when the system was inverted i.e. EphA2 LBD or ectodomains were immobilized via a histidine-tag to the sensor, and gH/gL was in solution (Fig. S7B).

Each of the 3 mutations introduced in EphA2 significantly reduced the binding as anticipated (Fig. 2D). The Q30N^gL^ mutation in the gL N-terminal segment also diminished binding, consistent with the presence of a carbohydrate at this position blocking the interactions with the strand D^EphA2^ and DE^EphA2^ loop. Introduction of the N-linked carbohydrate at residue N68^gL^ in its β2-β3 turn did not affect binding as expected, because of its location proximal to the binding site but in an exposed loop (Fig. 2B). The E52R^gH^/gL and E52A^gH^/gL variants resulted in weaker or absence of interactions with EphA2 ectodomains, respectively (Fig. 2D). These results demonstrated that the binding interface between EphA2 and gH/gL seen in the crystal is in agreement with the one mapped by measurements in solution.

### The gH/gL molecular mimicry of ephrin-A ligands

The EphA2 binding site for ephrin-A1 ligand and gH/gL largely overlap, with the former including a more extensive surface area and a larger number of contacts established by EphA2 β-strands D and E, CD and DE loops, as well as the LM loop (Fig. 3, Fig. S1 and S5). Comparative analyses of the gH/gL-EphA2 and ephrin-A1-EphA2 complexes further demonstrate that the structural elements employed by gH and gL resemble the ephrin-A1 ligand mode of binding to EphA2 receptor. The GH^ephrin-A1^ loop, which is the principal interaction region with the EphA2 receptor, occupies the same space as the gL ‘tail’, while the salt bridge established between the conserved R103^EphA2^ and E119^ephrin-A1^ is replaced at the same location by a salt bridge between R103^EphA2^ and E52^gH^ (Fig. 3). The conserved E52^gH^ and E119^ephrin-A1^ occupy equivalent position in respect to R103^EphA2^, but the chain segments carrying the conserved E52^gH^ and E119^ephrin-A1^ run in opposite directions so that the following residues, F53^gH^ and F120^ephrin-A1^, do not superpose. Both F53^gH^ and F120^ephrin-A1^ are engaged in π-π stacking interactions with F108 ^ephrin-A1^ and H53^gL^, respectively; in addition, the F108^EphA2^ establishes π-π interactions with Y21^gL^ or F111^ephrin-A1^, indicating a common mechanism for stabilization of the loop that presents the critically important glutamic acid residue for interactions with EphA2.

**Figure 3:**
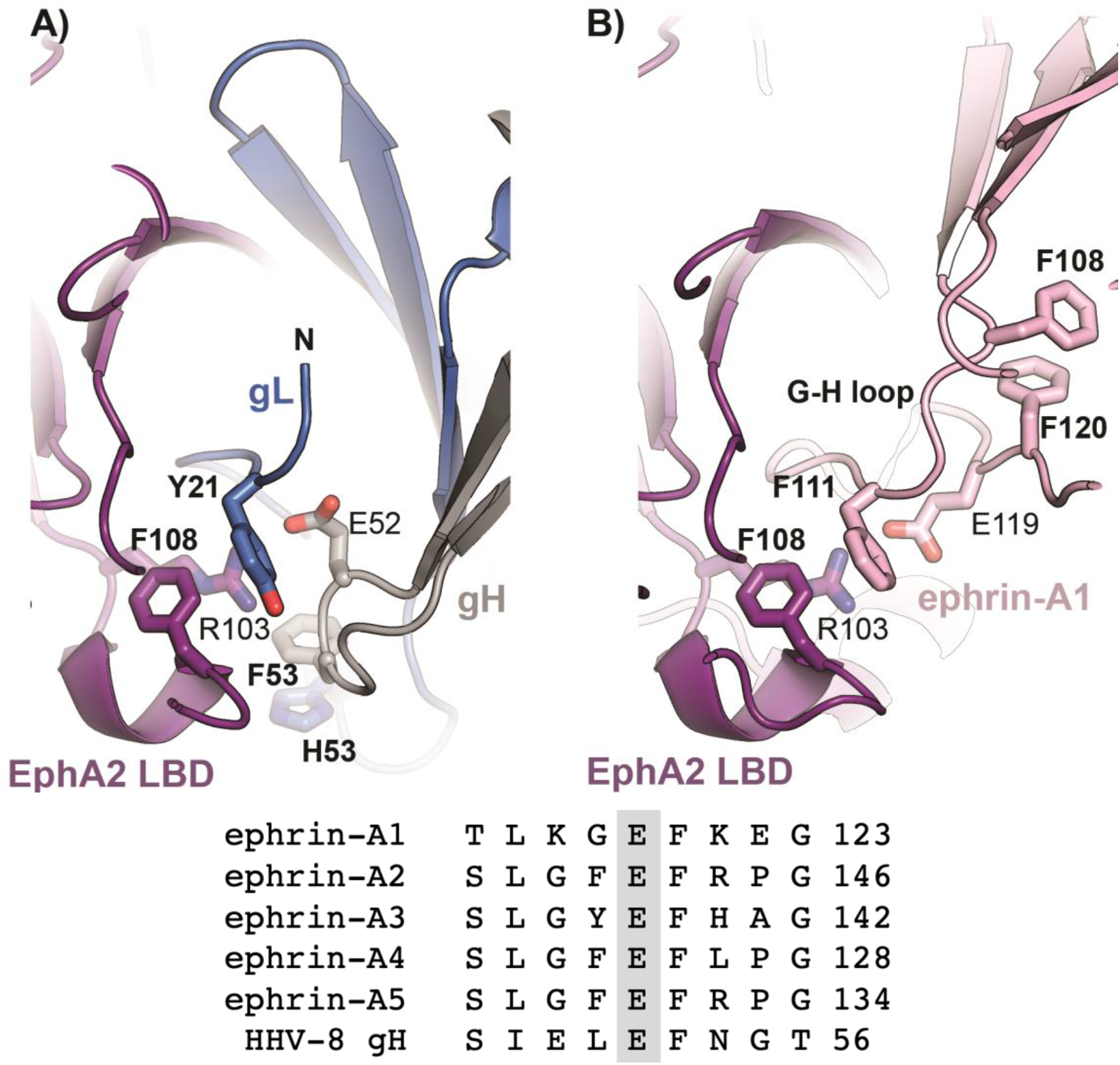
Structural mimicry between HHV-8 gH/gL and ephrin ligands. The EphA2 LBD from the EphA2 LBD - ephrin-A1 complex structure (PDB 3HEI) was superimposed onto the EphA2 LBD from our complex. The same coloring scheme for gH/gL and EphA2 is applied as in the previous figures, with the GH^ephrin-A1^ loop shown in pink. For clarity, only the elements participating in the interactions are shown. The E119 ^ephrin-A1^ is indicated. Sequence alignment of a GH loop segment of ephrin-A ligands, and the HHV-8 gH sequence is displayed to highlight the conservation of the glutamic acid that forms SBs with the EphA2^R103^ (E52^gH^ and E119^ephrin-A1^).

### HHV-8 gH/gL induces constitutive EphA2 dimerization on the cell surface

Since binding of ephrin ligands to Eph receptors induces formation of higher-order receptor oligomers, we sought to determine if gH/gL alters EphA2 interactions at the cell surface in a similar fashion. The method we applied was developed to probe the stability and association (stoichiometry) of protein complexes in cell membranes, and is referred to as FSI-FRET (Fully Quantified Spectral Imaging (FSI) Förster Resonance Energy Transfer (FRET)) ^53^. The measurements are carried out on the membranes of live cells containing the proteins of interest tagged with donor or acceptor fluorescent probes at the intracellular end. The lateral interactions of EphA2 molecules in the absence and presence of various ligands were already investigated using this approach ^47^.

We performed FSI-FRET measurements in HEK293T cells co-transfected with EphA2-mTurquoise (donor probe) and EphA2-eYFP (acceptor probe). Recombinant HHV-8 gH/gL was added at the final concentration of 200 nM, significantly exceeding the apparent subnanomolar Kd value (Fig. 2C), thus ensuring that all the EphA2 molecules were occupied by gH/gL. The measured FRET efficiencies (corrected for “proximity FRET” as discussed in MM (equation 1), and the concentration of donor-tagged and acceptor-tagged EphA2 molecules) were used to construct dimerization curves by fitting with a monomer-dimer equilibrium model ^53^ (the raw FRET data is shown in Fig. S8). Dimer formation is characterized by two parameters: the two-dimensional dissociation constant, *K_diss_*, and the structural parameter “Intrinsic FRET”, 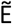. The *K_diss_* is a measure of the dimerization propensity of EphA2 at the plasma membrane. The Intrinsic FRET is the FRET efficiency in an EphA2 dimer with a donor and an acceptor, which depends on the positioning of the fluorescent proteins (attached to the C-terminus of the intracellular domain) of the EphA2 dimer. The Intrinsic FRET is strictly a structural parameter and therefore does not have any implications on the dimerization propensity of the full length EphA2 receptor [64, 65].

The dimerization curve calculated from the FRET data for EphA2 WT in the presence of gH/gL is shown in Fig. 4A and is compared to the data for EphA2 WT in the absence of ligand. The best-fit *K_diss_* and the best-fit Intrinsic FRET, determined from the FRET data, are presented in Table 1. As previously reported ^51^, EphA2 WT in the absence of ligand exists in monomer-dimer equilibrium with a *K_diss_* is 301 ± 67 receptors/μm^2^. We found EphA2 in the presence of gH/gL to be 100% dimeric (“constitutive dimer”) over the EphA2 concentration range observed in the experiments, precluding reliable measurement and calculation of the dissociation constant. In control experiments, we added soluble EphA2 LBD and also precomplexed gH/gL with EphA2 LBD (gH/gL-LBD) (Fig. 4B, C). As anticipated, soluble LBD had no effect on EphA2 dimerization and the effect of gH/gL was also abolished when precomplexed with EphA2 LBD, as the dimerization curves and the best-fit *K_diss_* value were indistinguishable from those determined in the absence of gH/gL (Table 1).

**Figure 4:**
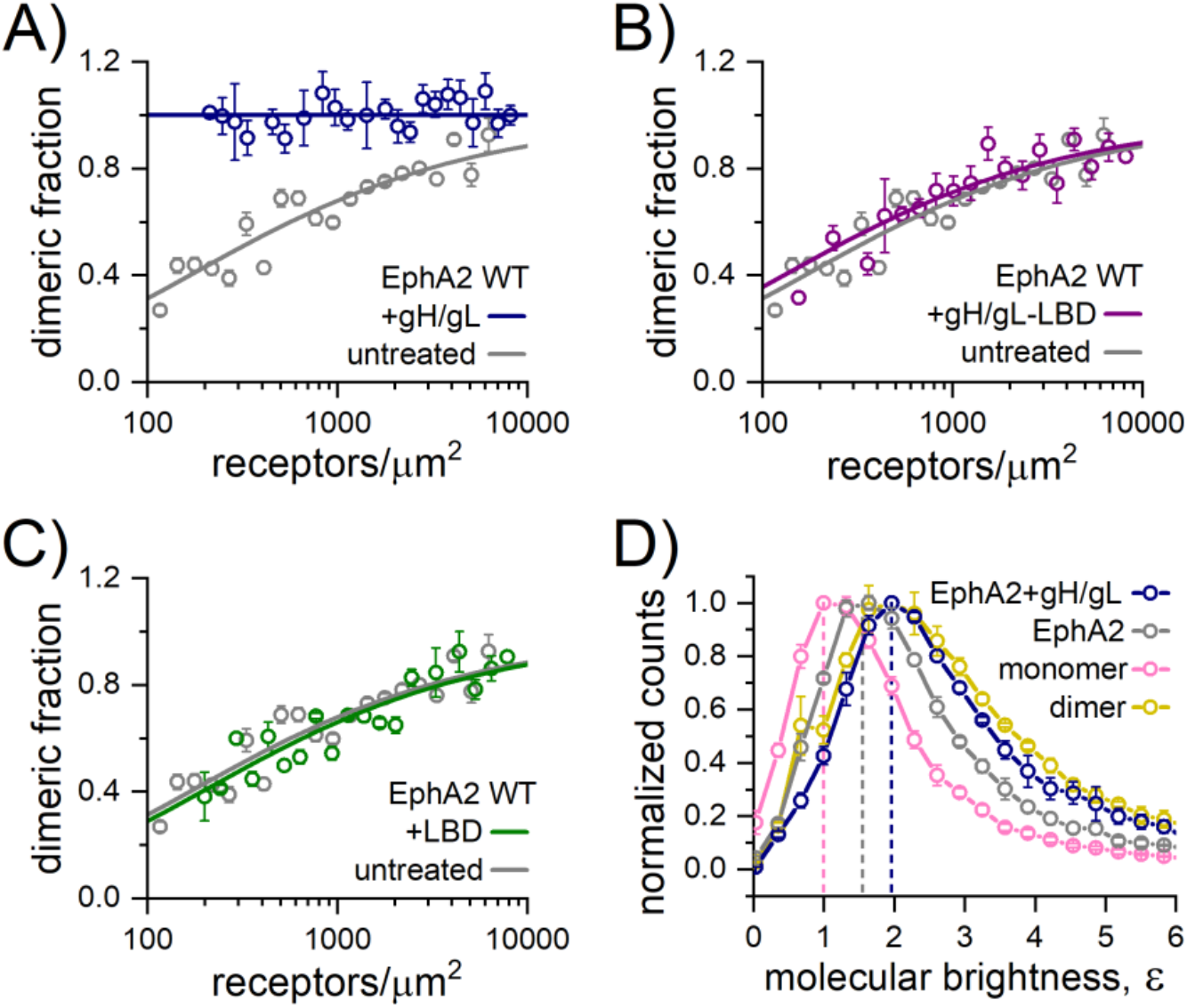
HHV-8 gH/gL induces constitutive EphA2 dimerization. The FSI-FRET data measured in HEK293T cells (Fig. S8) were fit to dimerization models to generate dimerization curves by plotting the calculated dimeric fraction as a function of the total EphA2 concentration. The binned dimeric fractions are shown along with the best-fit curve. The data measured for EphA2 WT in the presence of **(A)** 200 nM gH/gL, **(B)** 200 nM LBD, and **(C)** 200 nM gH/gL-LBD are compared to EphA2 WT data in the absence of ligand, which was previously reported [3]. Soluble gH/gL induces constitutive EphA2 dimerization, as evidenced by the dimeric fraction of 1 at all measured EphA2 concentrations. Little to no difference in the dimerization curves were observed when in the presence of LBD and precomplexed gH/gL-LBD compared to untreated EphA2 WT which suggests that EphA2 LBD blocks the effect of gH/gL on EphA2 dimerization. **(D)** Fluorescence Intensity Fluctuation (FIF) measurements in HEK293T cells reporting on EphA2 WT-eYFP oligomer size in the absence or presence of 200 nM gH/gL. Histograms of molecular brightness (*ε*), measured in small regions of the basolateral membrane over all measured receptor concentrations, are compared to the published FIF data for the monomer control (LAT) and dimer control (E-cadherin). The maximum of the histogram for EphA2 WT in the presence of gH/gL shifts to higher brightness than for EphA2 WT in the absence of ligand and is very similar to that of the E-cadherin dimer control, which suggests that EphA2 is a constitutive dimer in the presence of gH/gL, consistent with the FSI-FRET data.

**Table 1:**
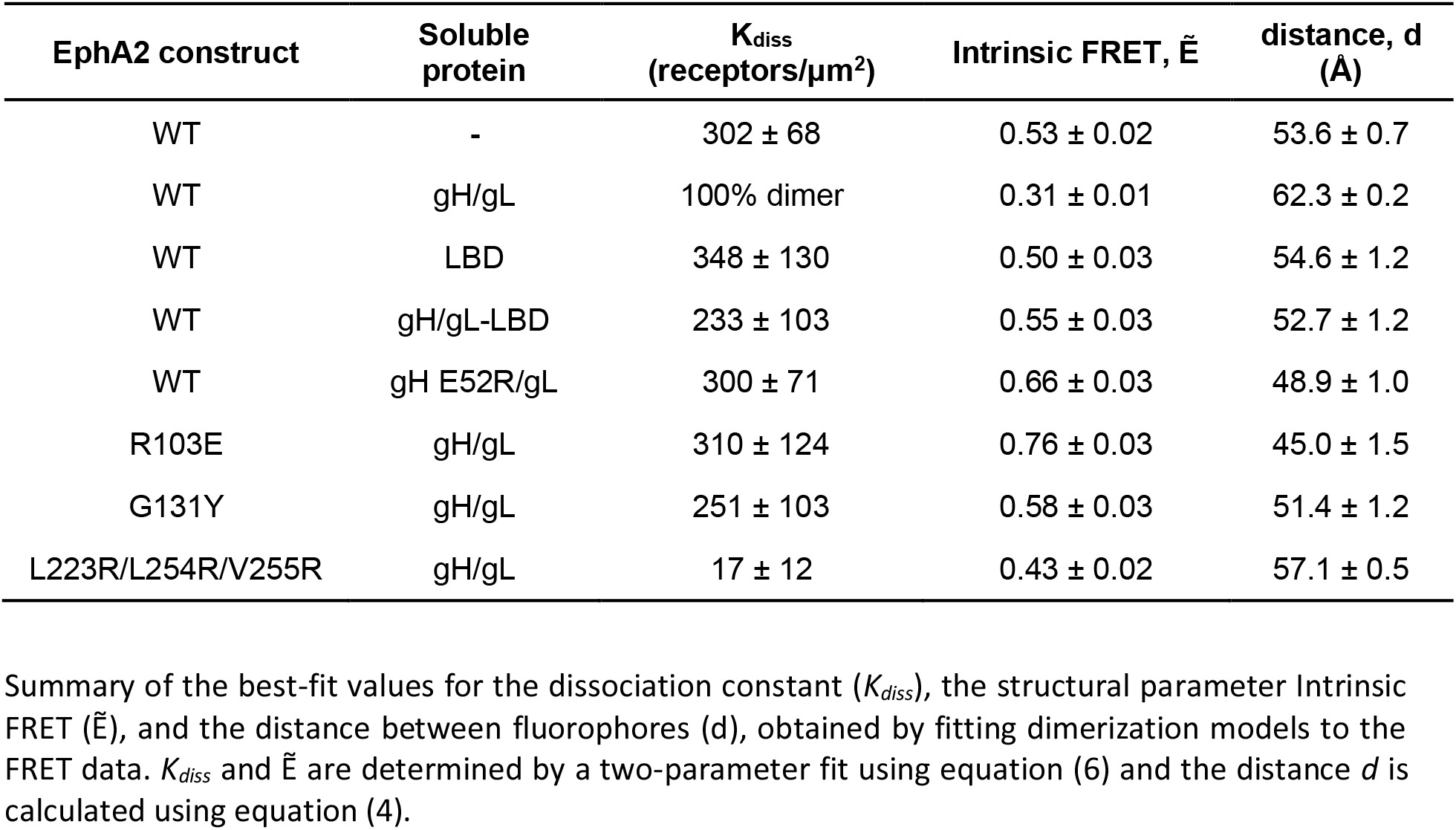
Sumary of the dimerization models fit to FRET data

| EphA2 construct | Soluble protein | $K_{diss}$<br>(receptors/$\mu\text{m}^2$) | Intrinsic FRET, $\tilde{E}$ | distance, $d$<br>(Å) |
| --- | --- | --- | --- | --- |
| WT | - | $302 \pm 68$ | $0.53 \pm 0.02$ | $53.6 \pm 0.7$ |
| WT | gH/gL | 100% dimer | $0.31 \pm 0.01$ | $62.3 \pm 0.2$ |
| WT | LBD | $348 \pm 130$ | $0.50 \pm 0.03$ | $54.6 \pm 1.2$ |
| WT | gH/gL-LBD | $233 \pm 103$ | $0.55 \pm 0.03$ | $52.7 \pm 1.2$ |
| WT | gH E52R/gL | $300 \pm 71$ | $0.66 \pm 0.03$ | $48.9 \pm 1.0$ |
| R103E | gH/gL | $310 \pm 124$ | $0.76 \pm 0.03$ | $45.0 \pm 1.5$ |
| G131Y | gH/gL | $251 \pm 103$ | $0.58 \pm 0.03$ | $51.4 \pm 1.2$ |
| L223R/L254R/V255R | gH/gL | $17 \pm 12$ | $0.43 \pm 0.02$ | $57.1 \pm 0.5$ |

Summary of the best-fit values for the dissociation constant ( $K_{diss}$ ), the structural parameter Intrinsic FRET ( $\tilde{E}$ ), and the distance between fluorophores ( $d$ ), obtained by fitting dimerization models to the FRET data. $K_{diss}$ and $\tilde{E}$ are determined by a two-parameter fit using equation (6) and the distance $d$ is calculated using equation (4).

Along with *K_diss_*, which measures the strength of the EphA2 association, these experiments give information about conformational changes that affect the relative disposition of the fluorescent proteins attached to the C-termini of EphA2, inside the cell. This information is contained in the structural parameter “Intrinsic FRET”. A lower Intrinsic FRET value is observed upon gH/gL binding, reflecting that the distance (*d*) between the fluorescent proteins is greater when gH/gL is bound to EphA2. Since the fluorescent proteins are attached to the C-termini of EphA2 via flexible linkers, this is a demonstration of a structural change in the EphA2 dimer induced by gH/gL binding, which is transmitted across the membrane to the intracellular domains, involving an apparent increase in the separation between the C-termini of EphA2. This implies that the conformation of the intracellular domains in the EphA2 dimer are altered in response to gH/gL binding. The presence of soluble EphA2 LBD and precomplexed gH/gL-LBD had no effect on the Intrinsic FRET values (Table 1).

Since FRET has limited utility in discerning the oligomer size, we used Fluorescence Intensity Fluctuation (FIF) to directly assess the oligomer size of EphA2 in the presence of gH/gL. FIF calculates molecular brightness of eYFP-tagged receptors in regions of the cell membrane. The molecular brightness, defined as the ratio of the variance of the fluorescence intensity within a membrane region to the mean fluorescence intensity in this region, is known to scale with the oligomer size ^61^. The cumulative (over all measured EphA2 concentrations) distributions of molecular brightness for EphA2 and EphA2 with gH/gL obtained from small sections of the plasma membrane in hundreds of cells are compared in Fig. 4D. Consistent with the fact that EphA2 exists in a monomer/dimer equilibrium in the absence of ligand ^51^, the EphA2 brightness distribution is between the distributions of LAT (Linker for Activation of T-cells, a monomer control) ^62^ and E-cadherin (a dimer control) ^63^. The FIF data for these controls have been published previously ^64^ and are shown here for comparison (Fig. 4D). We found that gH/gL shifts the maximum of the histogram to higher molecular brightness relative to EphA2 (untreated), such that it virtually overlaps with the dimeric E-cadherin distribution. Therefore, the FIF measurements indicate that EphA2 is a constitutive dimer in the presence of gH/gL, consistent with the FRET data. We see no indication for the formation of higher order oligomers, which would have resulted in a brightness distribution shifted to higher values than the ones measured for E-cadherin.

Taken together, the FRET and FIF data demonstrate that gH/gL significantly stabilizes EphA2 dimers but does not induce EphA2 oligomerization.

### Residues E52^gH^ and R103^EphA2^ are critical for EphA2 dimerization on cells

To test the importance of residue E52^gH^ for binding of gH/gL to EphA2 in native membranes, FSI-FRET experiments were also performed with the E52R^gH^/gL recombinant protein, which exhibited significantly reduced binding to the soluble EphA2 ectodomains in BLI experiments (Fig. 2D). The dimerization curve for EphA2 WT in the presence of E52R^gH^/gL is shown in Figure 5A and the fit parameters in Table 1. Constitutive EphA2 receptor dimerization was not observed as we did with gH/gL. Rather, EphA2 interactions were reduced to levels similar to the case of no ligand, indicating that the presence of E52R^gH^/gL did not result in EphA2 dimer stabilization. This is consistent with the finding that the binding of this E52R^gH^/gL variant to EphA2 was disrupted, and/or with the idea that bound E52R^gH^/gL did not enhance dimer stability. However, the measured Intrinsic FRET was slightly increased, as compared to no treatment, suggesting a decrease in the separation between the attached fluorescent proteins, and thus between the C-termini of EphA2. This effect could be due to structural perturbations in the EphA2 dimer in response to possible E52R^gH^/gL binding at the high E52R^gH^/gL (200nM) concentrations used, which could have propagated to the intracellular domain of EphA2.

**Figure 5:**
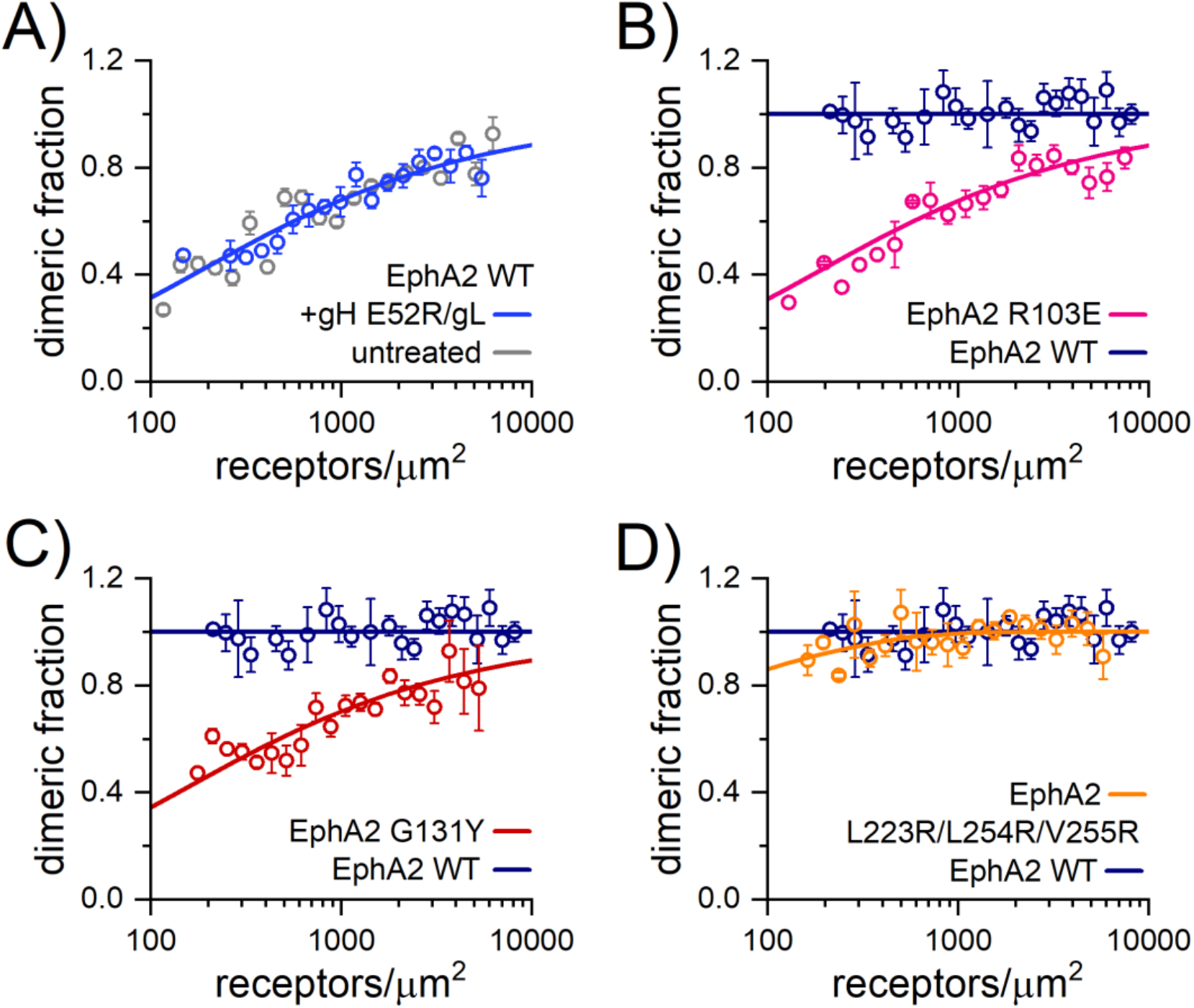
The gH/gL induced EphA2 dimers on cells engage the ‘dimerization’ interface. Dimerization curves calculated from the FSI-FRET data for **(A)** EphA2 WT in the presence of 200 nM gH E52R/gL mutant with mutation in EphA2 binding, and for the EphA2 mutants **(B)** R103E^EphA2^ mutant impaired in ligand binding, **(C)** G131Y^EphA2^ mutant with mutation in DIN, and **(D)** L223R/L254R/V255R^EphA2^ mutant with mutations in CIN. The data in A are compared to EphA2 WT data in the absence of ligand. The data in B-D were collected in the presence of 200 nM gH/gL and are compared to EphA2 WT in the presence of gH/gL (Fig. S8). No difference in the dimerization curve is observed with the mutated gH E52R/gL and thus does not induce constitutive EphA2 dimers as gH/gL does, which suggests impaired binding to EphA2. Large differences in the dimerization curves are observed for the R103E^EphA2^ and G131Y^EphA2^ mutants, but the effect of the triple L223R/L254R/V255R^EphA2^ mutation is modest. This data indicates that gH/gL-bound EphA2 dimers interact mainly via the DIN (where G131 is engaged) but not via the CIN (where L223/L254/V255 are engaged) and that R103^EphA2^ is important for gH/gL binding.

In addition, we sought to test the importance of the residue R103^EphA2^ for binding to gH/gL using the FSI-FRET method. The dimerization curve when the cells were transfected with EphA2 harboring the R103E mutation in the gH/gL binding site, in the presence of saturating gH/gL concentrations, is shown in Fig. 5B, and the fit parameters are shown in Table 1. The dimerization propensity for R103E^EphA2^ variant in the presence of gH/gL is the same as for EphA2 in the absence of ligand (Table 1), indicating that either gH/gL binding to the R103E^EphA2^ mutant is disrupted, as also seen in the BLI experiments, and/or that binding did not lead to dimer stabilization. These data further corroborate our findings that R103^EphA2^ plays an essential role in gH/gL binding. Here again, we observed an increase in the Intrinsic FRET, which indicates that the fluorescent proteins are in closer proximity, as compared to EphA2 WT in the absence of ligand (Table 1). Similar to the behavior of the E52R^gH^ /gL variant, this effect could be a consequence of R103A^EphA2^ binding to gH/gL, at the high gH/gL concentrations used, that would be transmitted to the EphA2 intracellular domains.

### HHV-8 gH/gL induced EphA2 dimers on cell surface are stabilized via the ‘dimerization’ interface

We showed that when bound to gH/gL, EphA2 is a constitutive dimer. To determine if the EphA2 dimers form via one of the already described interaction surfaces - the dimerization (DIN) or clustering interface (CIN), as reported previously ^47^ (Fig. S2), we transfected HEK293T cells with the EphA2 variants with perturbed dimerization (G131Y) or clustering (L223R/L254R/V255R) interfaces and treated them with soluble gH/gL ectodomains. The binding of these variants to gH/gL in solution, as measured by BLI, was not affected by the mutations, all of which reside outside of the gH/gL binding site (Fig. S2). The dimerization curves and FRET efficiencies for these mutants in the presence of gH/gL are shown in Fig. 5 and Fig. S8, respectively, with the fit parameters listed in Table 1. We observed a significant decrease in the dimerization due to the G131Y^EphA2^ mutation (Table 1; Fig. 5C), and a small effect due to the L223R/L254R/V255R^EphA2^ mutations (Table 1; Fig. 5D). Thus, G131^EphA2^ plays an important role in the stabilization of EphA2 dimers bound to gH/gL, implying that the EphA2 dimers are formed via DIN. This also supports our finding that gH/gL induces EphA2 dimers and not higher order oligomers, as these oligomers are known to engage both the DIN and the CIN (Fig. S2).

### HHV-8 gH/gL binding to EphA2 expressed on cells induces cell contraction

The interactions between Eph receptors and ephrin ligands is known to stimulate cell contraction, a signaling response that plays a role in developmental processes including axon guidance and tissue patterning ^65^. To test if the recombinant gH/gL proteins induce similar effects, we performed the assays in HEK293T cells, which express very low amounts of EphA2 (generally below the Western blot detection limit^47^). Therefore, we generated a HEK293T cell line that stably expressed EphA2, and measured cell contraction induced by gH/gL and the E52R^gH^/gL variant that bound weakly to EphA2 (Fig. 2D). The EphA2 LBD, alone or precomplexed with gH/gL (gH/gL-LBD) was used as negative control, and dimeric ephrin-A1-Fc as a positive control (described in the MM section). Untransfected HEK293T cells were treated with gH/gL in control experiments. The cells were fixed with paraformaldehyde (PFA), permeabilized, stained for actin and imaged (representative images are displayed in Fig. 6A). Histograms showing the mean cell area measured for each condition show that EphA2-HEK293T cells stimulated with gH/gL were significantly smaller in size compared to every other condition except for cells stimulated with dimeric ephrin-A1-Fc (Fig. 6B). This demonstrates that gH/gL triggered downstream signaling through EphA2, reminiscent to the ephrin-A1 induced signaling. Notably, untransfected HEK293T cells stimulated with gH/gL were also smaller than cells that were not stimulated, but not to the same extent as cells that stably express low levels of EphA2. When stimulated with E52R^gH^/gL, EphA2-HEK293T cells were slightly smaller than untreated cells, but not to the same extent as gH/gL-stimulated cells. The mean cell area determined for EphA2-HEK293T cells in response to E52R^gH^/gL was not statistically different from the area of untransfected HEK293T cells stimulated with gH/gL. Little to no differences in cell area are observed in EphA2-HEK293T cells upon incubation with EphA2 LBD or with the preformed gH/gL-LBD complex.

**Figure 6:**
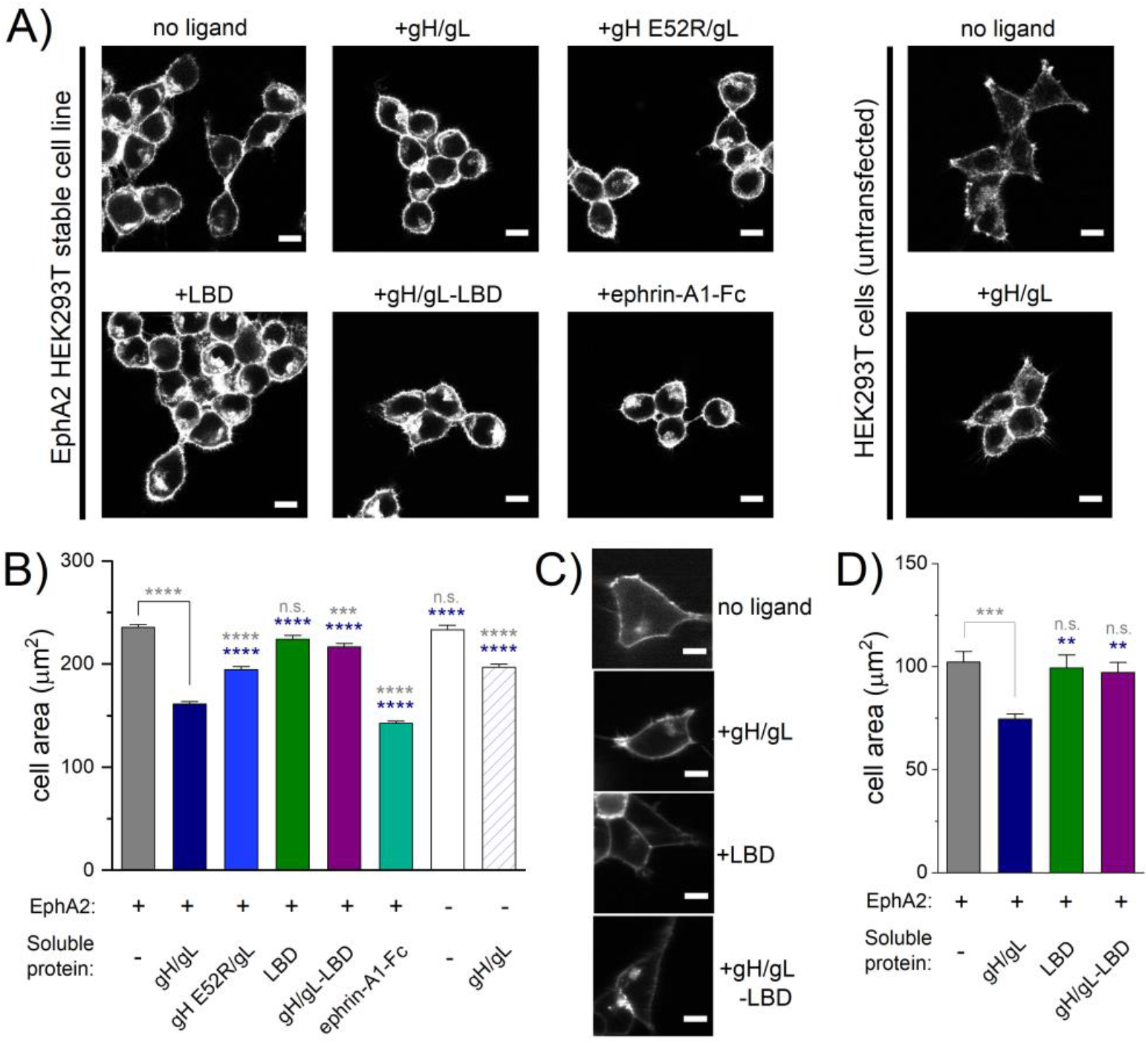
HHV-8 gH/gL stimulates EphA2-induced cell contraction. **A)** Images of fixed HEK293T cells stained for actin with rhodamine-conjugated phalloidin. The six images on the left were collected with HEK293T cells stably expressing EphA2 WT-eYFP and the two images on the right with HEK293T cells which do not express EphA2. Cells were stimulated with PBS (no ligand), 200 nM gH/gL, 200 nM gH E52R/gL, 200 nM LBD, 200 nM gH/gL-LBD complex, or 500 ng/mL ephrin-A1-Fc for 10 min prior to fixing with paraformaldehyde. Scale bar is 10 μm. **B)** Histograms of the average cell areas and the standard errors determined from the images of fixed cells shown in panel A. In the presence of saturating gH/gL concentrations, the EphA2-eYFP-expressing cells are significantly smaller in size compared to the cases of no ligand, +gH E52R/gL, +LBD, and +gH/gL-LBD, but are larger than cells stimulated with ephrin-A1-Fc. Untransfected HEK293T cells experience a slight decrease in average cell area in the presence of gH/gL but not to the same extent as cells expressing EphA2. Statistical significance was determined by a one-way ANOVA and a Tukey’s multiple comparison using the GraphPad Prism software (P < 0.0001 = ****, P < 0.001 = ***, P < 0.01 = **, P < 0.1 = *, P ≥ 0.1 = n.s.). Statistics results in grey are compared to EphA2 no ligand and those in navy are compared to EphA2 + gH/gL. **C)** Images of live HEK293T cells transiently transfected with EphA2 WT-eYFP in the absence or presence of 200 nM gH/gL, 200 nM LBD, or 200 nM gH/gL-LBD. Scale bar is 10 μm. **D)** Histograms showing the average cell areas and the standard errors determined from the live cell images shown in panel C. In the presence of saturating gH/gL concentrations, the EphA2-eYFP-expressing cells are smaller in size compared to cells with no ligand, with LBD, and with gH/gL-LBD. Statistical significance was determined by a one-way ANOVA and a Tukey’s multiple comparison using the GraphPad Prism software (P < 0.0001 = ****, P < 0.001 = ***, P < 0.01 = **, P < 0.1 = *, P ≥ 0.1 = n.s.). Statistics results shown in gray when comparison is made compared to unliganded EphA2, and those in navy in comparison to EphA2 + gH/gL.

To corroborate our findings in the fixed cells and exclude possible artifacts induced by PFA fixation, we performed a cell contraction assay without fixation using live HEK293T cells transiently transfected with full-length EphA2-eYFP. Soluble recombinant ectodomains of gH/gL, EphA2 LBD, or the gH/gL-LBD complex were added to the media. As in the fixed cell contraction assay, significant live cell area reduction was observed only when free gH/gL was added (Fig. 6C, D). These studies confirm that gH/gL mimics ephrin-A1 binding and signaling through EphA2.

## Discussion

The structure of the HHV-8 gH/gL complex bound to EphA2 LBD revealed two major gL elements important for binding - the N-terminal segment (res 22-30) and strands β2 and β3, the latter forming a mixed β-sheet with the EphA2 LBD (Fig. 2A). The gL residues form extensive van der Waals contacts and 14 hydrogen bonds with the LBD in total (Table S2A). Sequence alignment of gL from the gammaherpesvirus family shows preference for hydrophobic residues (Ala, Ile, Val) in the N-terminal segment (Fig. S10) consistent with the constraints imposed by packing of these side chains within the ‘channel’ formed by the hydrophobic residues from the EphA2 and gL β strands (‘roof’) (Fig. 2). In a cell-cell fusion assay, Su *et al.* reported that gH/gL from other gammaherpesvirus genera, the bovine Alcelaphine gammaherpesvirus 1 (AIHV-1) from the *Macavirus* genus, and Equid gammaherpesvirus 2 (EHV-2) from the *Percavirus* genus, bind to human EphA2 to trigger fusion, and suggested a potential for the spillover of animal herpesviruses to humans ^52^. They also demonstrated that this was not the case for the gH/gL from murid herpesvirus 4 (MHV68), which with HHV-8 belongs to the *Rhadinovirus* genus. We performed comparative sequence analyses and found that the N-terminal segment of MHV68 gL contains a number of charged residues (NH_3_^+^-<u>K</u>ILP<u>K</u>HCC…), which could preclude it from fitting into the human EphA2 binding ‘channel’. The same is true for gL from another rodent herpesvirus, cricetid gammaherpesvirus 2, whose N-terminus (NH_3_^+^-IIGSFL<u>A</u>RPCC) also contains a charged residue (Fig. S10), and we predict that this gH/gL complex would have weak binding affinity for human EphA2 as well. We therefore propose that the amino acid composition of the N-terminal gL segment can serve as a predictor for the potential for binding to human EphA2, and that the presence of charged or polar amino acids would weaken or abrogate the binding.

The C-terminal gL segment (residues 129-167) was not resolved in the structure, likely because the residues reside within a flexible region that seems to point away from the complex and is not involved in contacts neither with gH nor EphA2. This finding is consistent with the co-immunoprecipitation experiments done with HHV-8 gH co-expressed with the gLΔ135-164 variant, which could still form a complex with gH and EphA2 ^66^. This segment is absent from the EBV gL (Fig. S10) indicating a possibly virus-specific function.

Based on a series of BLI measurements we determined the Kd for WT HHV-8 gH/gL and EphA2 ectodomains to be in subnanomolar range. Decreasing the pH to 5.5, the endosomal pH at which the viral and endosomal membranes fuse to release the HHV-8 capsids into the cytoplasm ^67^, did not affect the Kd (Fig S7A), suggesting that gH/gL dissociation from EphA2 is not required for fusion. This would imply that gH/gL bound to EphA2 could still activate gB, or that there is a fraction of unliganded gH/gL that could interact with gB. The subnanomolar Kd value was obtained when the entire EphA2 ectodomain instead of the LBD alone was used, demonstrating that the EphA2 binding site for gH/gL resides in the LBD and that gH/gL does not interact with the downstream EphA2 domains. Higher dissociation constants for HHV-8 gH/gL and EphA2 of 3 or 9nM (depending on the orientation of the molecules) and 16nM were reported by two other groups, respectively ^52,60^. The discrepancies might be due to the overestimation of the gH/gL concentration, which was purified via a tag on gH, resulting in the protein preparation that contained gH/gL and free gH ^60^. Su *et al.* measured the affinities by SPR with the gH/gL immobilized to the chips by chemical coupling, procedure that modifies the protein surface and thus likely the binding site in a fraction of gH/gL ^52^. We designed the experiment to be able to separate the gH/gL complex from free gH by purifying the complex via the affinity tag on gL. The gH/gL was then immobilized to the sensors via the C-terminal affinity tag on gH, which is distal to the EphA2 binding site (Fig. S6). We therefore think that our data represent so far the most accurate quantification of the strength of gH/gL EphA2 interactions. The subnanomolar Kd for the WT proteins were obtained also in an inverted system i.e. when the EphA2 ectodomain or LBD were immobilized via affinity tag on its C-terminus and gH/gL was added as analyte (Fig S7B).

The residues of gH implicated in interactions with Eph receptors had been identified previously in a mutagenesis study ^66^. In particular, E52^gH^ and F53^gH^ in HHV-8, and the equivalent residues - E54^gH^ and F55^gH^ in rhesus RRV gH - had been found essential for EphA2 binding *in vitro*^24^. These residues are well conserved in gH of gamma-herpesviruses ^24^. The recombinant HHV-8 and RRV viruses carrying the mutations in this gH region were still able to attach to cells and infect them, although with one order of magnitude reduction in efficiency, by being re-targeted to other cellular attachment factors and receptors. We noticed that the gH β-turn (SIEL<u>EF</u>NGT) including E52^gH^ and F53^gH^ (underlined), carries resemblance to the motif located within a GH^ephrin-A1^ loop that is the main interaction structural element inserting into the EphA2 cavity (Fig. 3). A conserved glutamic acid residue in the same GH^ephrin-A1^ loop (E119^ephrin-A1^) forms salt bridge with the conserved R103 in the GH^EphA2^ loop (Fig. 3B, Fig. S1), which is the most important residue for ephrin binding, as its mutation to glutamic residue entirely abolished the interaction ^43^. In our structure the R103^EphA2^ forms 2 hydrogen bonds and 1 salt bridge with the E52^gH^, which is substantially less than the gL contribution (14 HB and extensive buried surface area) (Table S2). In EBV gH/gL the equivalent residue E30^gH^ is located away from the EphA2 binding interface due to a different organization of the N-terminal part of gH compared to the HHV-8 gH, possibly contributing to the reported weaker affinity with a Kd in the μM range ^52^ (the EBV gH and EphA2 in fact do not form any contacts). To determine to what extent the HHV-8 E52^gH^ influences interactions with EphA2, we performed biophysical (gH/gL and EphA2 ectodomains in solution) and cellular assays (EphA2 full-length membrane bound and gH/gL ectodomains in solution). We produced recombinant gH/gL variants carrying the mutations E52R^gH^ or E52A^gH^ and showed that both had reduced binding to EphA2 ectodomains in solution (Fig. 2D). When the cells expressing full-length EphA2 were treated with the recombinant HHV-8 E52R^gH^/gL, the changes reported for ephrin ligands ^68^ and induced by the WT gH/gL, such as increased EphA2 dimerization and cell contraction (Fig. 4A, Fig. 6), were not observed. Our data therefore demonstrate that the E52^gH^ is an important molecular determinant for EphA2 binding, further highlighting the gH/gL mimicry with ephrin ligands. While the single structural element of ephrins, the GH loop, engages in polar as well as hydrophobic interactions with EphA2, in HHV-8 gH/gL the same contributions are made by E52^gH^ and the N-terminal segment of gL.

EphA2 receptors are present on the cell surface in a dynamic monomer-dimer equilibrium, which shifts towards dimers and dimer-made oligomers as a function of ephrin ligand concentration ^18,69^. EphA2 signaling oligomers can be stabilized via one of the two opposite interfaces - the DIN and the CIN. In the absence of ligand, the EphA2 dimers are stabilized via CIN ^47^. The initial formation of a tetramer – a pair of EphA2 receptor-ligand complexes - is thought to be driven by the ephrins that behave as bivalent ligands interacting with their EphA2 partner molecule through the high-affinity site, and through a secondary site with the other EphA2 molecule from the same tetramer, holding together the whole assembly (Fig. S2). In this arrangement the EphA2 dimers were reported to be stabilized via the GH^Epha2^ loops (DIN), while the larger clusters that form as the ligand concentration increases are formed via contacts between adjacent CRD interfaces (CIN) (Fig. S2) ^44,45^. In other words, the association via DIN results in EphA2 dimers, while CIN interactions can result in EphA2 dimers or larger oligomers. (Fig. S2). Our data indicate that once the gH/gL binds, a structural transition is induced and EphA2 dimers are formed via DIN. To explore why EphA2 dimers and not larger oligomers were observed in our experiments, we constructed a model in which unliganded EphA2 ectodomains (PDB 2X10) ^29^ were superimposed onto the EphA2 LBD bound to the gH/gL, preserving the packing of the LBD-gH/gL we observed in the crystal (Fig. S9). This theoretical model reflects the putative arrangement EphA2 receptors expressed on cells would adopt upon binding to gH/gL, and indicates that the CINs would be too far apart to mediate EphA2 clustering, corroborating our ‘gH/gL induced EphA2 dimer’ finding.

The EphA2 dimers we observe in the crystal structure and in FSI-FRET studies, formed via the DIN, would presume ligand-driven stabilization resembling the formation of the Eph-ephrin tetramers described above. Analyses of the contacts in our structure show however the gH/gL molecules do not make contacts with the second EphA2 from the DIN-stabilized tetramer, but rather with an EphA2 from the neighboring EphA2 tetramer (Fig. S9). This poses question of how the gH/gL-induced EphA2 dimers are stabilized, and why CIN-stabilized EphA2 dimers bound to gH/gL are not observed on the cell-surface instead. We analyzed the DINs in unliganded EphA2 receptor and in complex with ephrin ligands and HHV-8 gH/gL. When EphA2 is bound to an ephrin ligand, the DIN is stabilized by increased buried surface area and more hydrogen bonds compared to unliganded EphA2 (Table S3). The values obtained for gH/gL induced EphA2 dimerization (16 HBs and 693 A^2^ interface) are comparable to the stabilization of the same interface when bivalent ephrin ligands are bound, indicating that binding of gH/gL to EphA2 LBD stabilizes the DIN sufficiently even though gH/gL does not seem to behave as a bivalent ligand bridging the two EphA2-gH/gL together. Our model would indicate, although this remains speculative, that gH/gL would preferentially bind to the monomeric EphA2 receptors on cells, shifting the equilibrium between the CIN-stabilized, unliganded EphA2 dimers to monomeric EphA2, which once bound to gH/gL would be stabilized via the DIN.

Ephrin ligands are expressed on cells as membrane-bound monomers, and Eph receptor clustering and activation at high level are *in vitro* typically induced by addition of soluble dimeric or pre-clustered ephrin ectodomains that mimic their ephrin high concentration when expressed at the cell surface. Ephrin-A1 ligand was also found as a soluble molecule, being released from cancerous cells by cleavage by cellular proteases. Such soluble, monomeric ephrin (m-ephrin) was demonstrated to be a functional ligand, able to activate the EphA2 receptor by inducing tyrosine phosphorylation ^70^, internalization of EphA2 and cell retraction, overall decreasing the cellular oncogenic potential ^71^. These beneficial cellular responses are characterized as the outcome of the so-called canonical Eph receptor activation ^72^. The structural and functional mimicry of HHV-8 gH/gL and ephrin-A1 would suggest that the HHV-8 interactions with EphA2 trigger canonical signaling pathways, consistent with the observed increased endocytosis and overall EphA2 phosphorylation upon virus binding ^20^, as well as gH/gL induced cell contraction that we report in Fig. 6. Chen *et al.* reported a different outcome when ephrin-A1 ligands were presented to EphA2-expressing cells in a polarized manner; the Src mediated signaling was activated instead, promoting cell motility and malignancy via phosphorylation of serine and not tyrosine residues (the so-called non-canonical EphA2 activation) ^48^. In that light it is interesting that HHV-8 binding to fibroblasts was also reported to result in Src recruitment by androgen receptor, a steroid-activated transcription factor that interacts with the intracellular domain of EphA2. Src activation in this case led to phosphorylation of the S897^EphA2^ in the intracellular EphA2 domain i.e. activation of the non-canonical pathway, which was a prerequisite for HHV-8 infection ^73^. These conflicting observations raise the question of what type of signaling HHV-8 gH/gL activates to enter the cells. The canonical and non-canonical pathways were thought to be mutually exclusive, but Barquilla *et al.* recently reported that the two can co-exists and that the EphA2 canonical signaling can be rewired in prostate cancer cells by androgenic receptors leading to the phosphorylation of the same S897^EphA2^ residue ^30^ implicated in HHV-8 entry ^73^. The authors proposed that this EphA2 reprogramming could have implications for the disease progression. It will be important to discern if a similar interplay of the two seemingly antagonistic EphA2 pathways exist in cells infected with HHV-8. It is also possible that there is a temporal regulation and that different EphA2 signaling pathways are activated during the primary infection (canonical, which would stimulate virus internalization via endocytosis) and reactivation (non-canonical, which would increase cellular oncogenic potential).

In this report we present the structure of the HHV-8 gH/gL bound to EphA2, and show that the gH/gL induces dimerization of EphA2 expressed by cells as well as morphological changes at cellular level, resembling the action mode of ephrin ligands. It is possible that the membrane anchored gH/gL at the virion surface could induce EphA2 oligomerization into even larger aggregates, in particular if membrane regions with higher local gH/gL concentration exist in virions, emulating the conditions of high ephrin ligand concentration. What is clear however is that at the mechanistic level HHV-8 gH/gL and ephrin ligands induce formation of the same EphA2 dimers, and that these dimers are already functional leading to cytoskeletal rearrangements and cell contraction. How the structural changes that gH/gL binding initiates are transmitted to the EphA2 intracellular domain, and which types of signaling cascades are elicited and when during the virus life cycle are the questions that need to be addressed next.

## Material and methods

### Protein expression and purification for crystallization

#### HHV8 gH/gL

The segments coding the ectodomain of HHV8 gH (residues 26 to 704) and the entire gL (residues 21 - 167) were each cloned from the already described plasmids ^20^ into the pT350 vector ^74^ for expression in Schneider S2 *Drosophila* cells (S2 cells). The Cys58 on gL was predicted to be unpaired based on the sequence alignment with EBV gL and was mutated to Ser to prevent potential formation of disulphide-linked dimers. The pT350 vector contains inducible metallothionein promoter activated by divalent cations, the exogenous *Drosophila* Bip signal peptide (<u>MKLCILLAVVAFVGLSLG</u> *RS*), underlined, that drives protein secretion appended to the N-terminus of the protein upstream of the 5’ cloning site. The *RS* are vector residues that code for the BglII cloning site. A double strep tag (DST) for affinity purification at the C-terminus was added downstream of the 3’ cloning site. The sequence of the DST in our pT350 plasmid is FE<u>DDDDK</u>*AG***WSHPQFEK***GGGSGGGSGGGS***WSHPQFEK**. where DDDDK is the enterokinase cleavage site and the sequences in bold correspond to the two strep tags separated by a GS linker (italics). The residues FE come from the vector *BstBI* cloning site.

HHV-8 gH can be secreted without being bound to gL ^75^, and the complex purification via a gH tag resulted in a mixture of the gH/gL complex and free gH, which were difficult to separate on size exclusion chromatography (SEC) due to the small size difference (gH has a molecular weight of 75 kDa, and gL 20 kDa). This is why it was crucial for gH/gL purification to add the tag for affinity purification only on gL (Fig. 1A). Thus, the gL was cloned to contain the DST at the C-terminus, and gH has a stop codon before the DST. The resulting expression plasmids were named gH^nt^pT350 and gLC58S^st^pT350, respectively, where ‘nt’ and ‘st’ stand for no tag and strep tag, respectively. Such choice was made to allow affinity purification of the gH/gL complex via gL.

It is worth noting that due to the design of the pT350 vector for cloning via restriction digestion the expressed proteins end up having two extra residues RS (*BglII* site) at the N-terminus. The gLC58S^st^pT350 and gH^nt^pT350 both contain the RS before the first authentic residues of the mature gL (Y21) and gH (L26).

The gH^nt^pT350 and gLC58S^st^pT350 plasmids were co-transfected in the S2 cells along with the plasmid encoding puromycin resistance, and the stably transfected cell lines were established by puromycin selection during 3-4 weeks following the previously established protocol ^76^. The expression was induced by addition of 0.5mM CuSO_4_, and the protein was purified from the supernatant 7-10 days post induction. After purification on Streptactin column (IBA Biosciences) and Superdex S200 16/60 GL column (GE life sciences), around 1 milligram of the gH/gL heterodimer was obtained per liter of cell culture (the same amount of the aggregated protein was present). SEC running buffer was 10 mM Tris, 50mM NaCl pH 8.0.

#### EphA2LBD

The EphA2 gene segment that encodes for residues 23-202 was amplified from the vector ^20^ and cloned into the pT350 vector for expression in S2 cells (EphA2LBD^st^pT350 construct). The expressed protein contained the DST at the C-terminus. Affinity and SEC purifications were done as described for gH/gL above. Around 10 milligrams of pure EphA2 LBD were obtained from 1 liter of cell culture.

#### Preparation and purification of the trimeric complex EphA2 LBD-HHV8 gH/gL for crystallization

HHV-8 gH has 14 predicted N-glycosylation sites, gL one and EphA2 LBD none. To increase the probability that the complex would crystallize, the gH/gL complex was enzymatically deglycosylated with recombinant Endoglycosydase D (endo-β-N-acetylglucosaminidase from *Streptococcus pneumoniae* i.e. EndoD ^77^), in 100mM sodium-citrate buffer pH 5.0, 150mM NaCl for 18h at 25°C. The ratio of protein to EndoD was 40 to 1 (w:w). To remove the EndoD and exchange the reaction buffer, the deglycosylated complex was purified on Superdex S200 column in 10 mM Tris, 50mM NaCl pH 8.0, and then mixed with the purified EphA2 LBD in 1:1.3 molar ratio (gH/gL: EphA2 LBD). The complex was incubated at 4°C for 1h to over-night, and then re-purified on Superdex S200 to separate the excess EphA2 LBD. The running buffer in all SEC purifications was 10mM Tris pH 8, 50mM NaCl. The trimeric complex was stable on SEC and presence of gH/gL and the EphA2 LBD in the complex peak was verified by SDS-PAGE gel analysis.

### Crystallization

The tertiary complex in 10 mM Tris pH 8, 50 mM NaCl was concentrated to 5.1 mg/ml in Vivaspin concentrators with the 10 kDa cut-off. Crystallization screening was performed at the Institut Pasteur Core facility for crystallization ^78^ by vapor diffusion, in sitting drops of 0.4 μl containing equal volumes of the protein and reservoir solution. The drops were dispensed in 96-well Greiner plates by a Mosquito robot (TTPLabtech, Melbourn, UK) and images were recorder by a Rock-Imager 1000 (Formulatrix, Bedford, MA, USA). Initially, crystals were found in numerous conditions, but none diffracted better than 5Å. To improve these crystals, Hampton additive screen HT (HR2-138) was set up next based on one of the initial hits, and crystals grown in 0.1M Na-malonate pH 5, 14.2% PEG 3350 in the presence of 14 mM adenosine-5’-triphosphate disodium salt hydrate diffracted to 2.7 Å. These crystals were transferred into the crystallization solution supplemented by 20% ethylene glycol as a cryo-protectant and flash-frozen in liquid nitrogen.

### Data collection and structure determination

Data collected at the Proxima 1 beamline at the French national synchrotron facility (SOLEIL, St Aubin, France) were indexed, integrated, scaled and merged using XDS^79^ and AIMLESS^80^. Molecular replacement was done with Phaser within Phenix ^81 82^ using as search models the EphA2 LBD structure (Protein Data Bank [PDB] accession number 3HEI) and the HHV8 gH/gL model that was generated in Phyre2 ^55^ based on the sequence similarity with the EBV gH/gL (PDB code 3PHF). The partial solution containing EphA2 LBD and parts of gH was obtained, and was extended by iterative rounds of model building (Autobuild ^83^ in Phenix and manual building in Coot^84^) and refinement using Buster^85^ and Phenix ^82^. The final model converged to R_work_/R_free_ of 0.22/0.24. The final map displayed clear electron density for residues 27-200 of EphA2 (with a break in the J helix region (residues 148-162) and GH loop (G111)), residues 21-128 for gL, and residues 35-696 gH with the exception of several short regions of poor density that precluded unambiguous placement of the polypeptide chain (Fig. 1A).

The crystallization conditions, crystal parameters, data statistics, and refinement parameters are shown in Table S1. Superpositions of structures and all structural figures were generated with PyMOL (version 1.3r1)^86^. The atomic coordinates and structure factors for trimeric complex EphA2 LBD-HHV8 gH/gL have been deposited in the RCSB Protein Data Bank with the PDB code 7B7N.

### Expression of HHV8 gH/gL and EphA2 variants in mammalian cells

To avoid lengthy selection of the stably transfected S2 cells expressing recombinant proteins (4-5 weeks), a panel of gH/gL and EphA2 variants to be tested in biophysical assays was ordered as synthetic genes (GenScript) cloned in pcDNA3.1 (+) vector for transient expression in mammalian Expi293 cells (5-7 days). The reason mammalian cells were not used for production of the proteins used for crystallizations is that gH/gL complex is heavily glycosylated and the complex sugars added by mammalian cells are typically detrimental for protein crystallization and cannot be enzymatically removed unlike the simple Endo-D and Endo-H sensitive carbohydrates added by insect cells.

The mammalian cell expression constructs for gH and gL (residues 21 – 167, and 26-704 respectively) contained at the N-termini an exogenous murine Ig κ-chain leader sequence that targets protein to secretory pathway (Coloma et al., 1992) (METDTLLLWVLLLWVPGSTG). Different affinity tags were added to their C-termini - for gH construct (gH^his^pcDNA3.1) an enterokinase cleavage site (underlined) flanked by two GS linkers (italics) was followed by an octa-his tag (bold) (*GS* <u>DDDDK</u> *SGS* **HHHHHHHH**), and for gL a DST (gL^st^pcDNA3.1; the DST sequence is the same as in the gLC58S^st^pT350 construct described above). The EphA2 variants contained the endogenous signal peptide at the N-terminus and ended at residue 534 followed by a DST tag.

### Biophysical analyses

#### SEC-MALS measurements

The complexes were formed by incubating gH^his^/gL^st^ and EphA2^st^, both produced in mammalian cells as described above, in 1:1.3 molar ratio for 30 minutes at 4°C (~100 μg of gH/gL were mixed with 80 μg of EphA2 ectodomains or 30 μg of the EphA2 LBD in PBS in total volume of 200 μl). Assembled complexes were injected into Superdex 200 10/300 GL column (GE life sciences) using 500 μl loop, and run in PBS at a flow rate of 0.4 ml/min. The samples passed through a Wyatt DAWN Heleos II EOS 18-angle laser photometer coupled to a Wyatt Optilab TrEX differential refractive index detector. Data were analyzed with Astra 6 software (Wyatt Technology Corp).

#### Biolayer interferometry measurements

Biolayer interferometry (BLI) was applied to determine the effect of gH/gL and EphA2 mutations on the complex dissociation constant (Kd). The measurements were carried out on an Octet RED384 instrument (ForteBio). Affinity and SEC purified gH^his^/gL^st^ produced in mammalian Expi293 cells was immobilized on Ni^2+^-NTA sensors (ForteBio) in PBS. The loaded and equilibrated biosensors were dipped into analyte solutions containing 250 nM to 1 nM EphA2 variants in PBS containing 0.2 mg/ml BSA (the assay buffer).

Association and dissociation were monitored for 250 and 500 seconds respectively. Sensor reference measurement was recorded from a sensor not loaded with gH/gL and dipped in PBS. Sample reference was recorded from a sensor loaded with gH/gL that was dipped in the assay buffer. Specific signals were calculated by double-referencing, that is, subtracting non-specific signals obtained for the sensor and sample references from the signals recorded for the gH/gL-loaded sensors dipped in EphA2 analyte solutions. Association and dissociation profiles, as well as steady-state signal versus concentration curves, were fitted assuming a 1:1 binding model.

### Cell culture used in FRET and contraction experiments

Human embryonic kidney (HEK) 293T cells were purchased from American Type Culture Collection (Manassas, VA; CRL-3216). The cells were cultured at 37 °C and 5% CO_2_ in Dulbecco’s Modified Eagle Medium (DMEM; Thermo Scientific; 31600-034) that contained 3.5 g/L D-glucose, 1.5 g/L sodium bicarbonate, and 10% fetal bovine serum (FBS; Sigma-Aldrich; F4135). The cells were passed up to 25 times and then discarded.

### Fixed cell contraction assay

HEK293T cells transiently transfected with WT EphA2-eYFP were selected with 1.6 mg/ml G-418 solution (Roche; 4727878001) for 12 days to generate a stable cell line. The concentration of G-418 was determined using a kill curve. HEK293T cells or EphA2 HEK293T stable cells were seeded (1 x 10^4^ cells/well) into 8-well tissue culture chambered coverglass slides (Thermo Scientific; 12565338) and cultured for 36 hours. The cells were then washed twice with serum-free, phenol red-free media and were serum starved for 12 hours overnight. The cells were washed twice with PBS, followed by treatment for 10 min at 37°C with PBS or 200 nM gH/gL, 200 nM gH E52R/gL, 200 nM LBD, 200 nM gH/gL-LBD, or 0.5 μg/ml ephrin-A1-Fc in PBS. The cells were then fixed in 4% paraformaldehyde in PBS for 15 min at 37°C, permeabilized in 0.1% Triton X-100 in PBS for 15 min 37°C, and incubated with blocking solution (5% FBS, 1% BSA in PBS) for 30 min at room temperature. Then, the cells were stained for actin using rhodamine-conjugated phalloidin (Thermo Fisher Scientific; R415) for 90 min at room temperature. The lyophilized phalloidin-rhodamine powder was reconstituted in DMSO to make a 40x stock solution and then diluted with blocking buffer to make a 1x solution which was added to the fixed cells. The cells were washed twice with PBS in between each step. Finally, starvation media was added to each well prior to imaging. Actin-stained fixed cells were imaged with a Leica TCS SP8 confocal microscope equipped with a HyD hybrid detector and a 63x objective. The measurements were performed with a 552 nm excitation diode laser at 0.5% power using the dsRed setting which measures the fluorescence between wavelengths of 562 and 700 nm. The scanning speed was at 200 Hz, the pixel depth at 12-bits, the zoom factor at 1, and the image size at 1024×1024 pixels. Cell area was determined using the ImageJ software (NIH, Bethesda, MD) by drawing a polygon around the membrane of the cells. Statistical significance was determined by one-way ANOVA followed by a Tukey’s test using the GraphPad Prism software.

### Live cell contraction assay

HEK293T cells were seeded, transfected with EphA2 WT-eYFP, and serum-starved overnight in the same manner as described in the FRET section. Ten minutes prior to imaging, the media was replaced with starvation media or starvation media containing 200 nM gH/gL, 200 nM LBD, or 200 nM gH/gL-LBD. Cells were imaged with a Zeiss Axio Observer Inverted two-photon microscope using a 63x objective at a wavelength of 960 nm to excite the eYFP fluorophore. Cell area and statistical significance was determined in the same manner used for the fixed cell contraction assay.

### FSI-FRET measurements and analysis

For FRET experiments, the cells were seeded in 35 mm glass bottom collagen-coated petri dishes (MatTek Corporation, MA) at a density of 2 x 10^5^ cells/dish and cultured for 24 hours. The cells were co-transfected with EphA2-mTurquoise (mTURQ, the donor) and EphA2-enhanced yellow fluorescent protein (eYFP, the acceptor) in pcDNA, as described ^87,88^, using 0.5-2 μg of total DNA and the Lipofectamine 3000 reagent (Invitrogen, CA). In control experiments, cells were transfected with either EphA2-mTURQ or EphA2-eYFP and used for calibration as described ^53^. Twelve hours following transfection, the cells were washed twice with serum-free, phenol red-free media and serum-starved in the same media for 12 hours overnight. Immediately before imaging, the starvation media was replaced with hypo-osmotic media (10% starvation media, 90% diH2O, 25 mM HEPES) to ‘unwrinkle’ the highly ruffled cell membrane under reversible conditions as described ^89^. The soluble proteins were premixed with the hypo-osmotic media before adding to the cells. Cells were incubated for 10 minutes and then imaged under these conditions for approximately 1 hour.

Spectral images of HEK293T cells under reversible osmotic conditions were obtained following published protocols ^53^ with a spectrally resolved two-photon microscope (Zeiss Inverted Axio Observer) equipped with line-scanning capabilities (OptiMis True Line Spectral Imaging system, Aurora Spectral Technologies, WI) ^90^. Fluorophores were excited by a mode-locked laser (MaiTai™, Spectra-Physics, Santa Clara, CA) that generates femtosecond pulses between wavelengths 690 nm to 1040 nm. Two images were collected for each cell: one at 840 nm to excite the donor and a second one at 960 nm to primarily excite the acceptor. Solutions of purified soluble fluorescent proteins (mTURQ and eYFP) at known concentrations were produced following a published protocol ^91^ and imaged at each of these excitation wavelengths. A linear fit generated from the pixel-level intensities of the solution standards was used to calibrate the effective three-dimensional protein concentration which can be converted into two-dimensional membrane protein concentrations in the cell membrane as described ^53^. The calibration curve along with the cell images were used to calculate the FRET efficiency and the concentration of donors and acceptors present in the cell membrane ^53^. Regions of the cell membrane not in contact with neighboring cells were selected and analyzed to avoid interactions with proteins on adjacent cells.

The measured FRET efficiencies (*E_app_*) were corrected for ‘proximity FRET’ (*E_prox_*) as described previously ^92^. The proximity FRET accounts for donor-tagged molecules and acceptor-tagged molecules coming into close enough proximity to observe a FRET signal (within 100 Å) but not interacting directly. The corrected FRET due to specific interactions between the labeled proteins is given by:

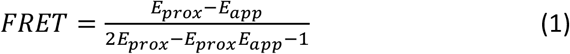

The corrected FRET depends on the fraction of membrane protein dimers, *f_D_*, and on the acceptor fraction, *x_A_*, according to:

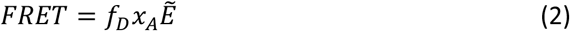

The ‘Intrinsic FRET’ (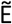) is a structural parameter that depends on the distance between and orientation of the two fluorophores in the dimer but is not dependent on the dimerization propensity ^92–94^.

When the dimeric fraction is 100% (*f_D_* = 1), the corrected FRET does not depend on the concentration of the labeled proteins and thus equation (2) can be simplified further:

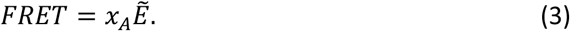

The dependence of the Intrinsic FRET on the distance between the fluorescent proteins in the dimer, *d*, is given by ^93^:

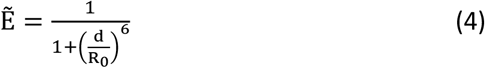

where *R_0_* is the Forster radius for the mTurquoise-eYFP FRET pair, 54.5 Å. Since the fluorescent proteins are attached to the C-terminus of the membrane proteins via long flexible linkers, we assume free rotation of the fluorescent proteins.

By rearranging equation (2), the dimeric fraction, *f_D_*, can be determined from the corrected FRET efficiencies and concentrations according to:

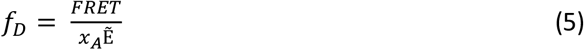

In the case of dimers, the following equation is used to determine the two unknowns *K_diss_* and 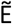 as described previously ^53^:

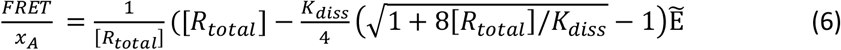

### Fluorescence intensity fluctuations (FIF) spectroscopy measurements and analysis

HEK293T cells were seeded as described in the FRET section at a density of 4 x 10^5^ cells/dish. The cells were transiently transfected 24 hours later with 1 μg of EphA2-eYFP using Lipofectamine 3000 and then washed and serum starved twelve hours later. The starvation media was replaced with a 75% hypo-osmotic media (25% starvation media, 75% diH2O, 25 mM HEPES) containing 200 nM gH/gL prior to imaging. A TCS SP8 confocal microscope (Leica) using the photon counting capabilities of the HyD hybrid detector was used to collect images of the cell basolateral membranes. The measurements were performed with a 488 nm excitation diode laser and the emission spectra of YFP were collected from 520-580 nm. The scanning speed was at 20Hz, the pixel depth at 12-bits, the zoom factor at 2, and the image size at 1024×1024.

The cell images were analyzed using the FIF software described in ^61^. The software performed segmentation of the basolateral membrane into 15×15 pixel regions of interest. Each cell is outlined by researcher prior to segmentation. The segmented data was analyzed using the brightness and concentration calculator in the FIF software ^61^. The molecular brightness, ε, was calculated according to:

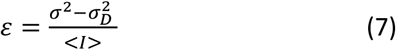

where *σ*^2^ is the variance of fluorescence across segments, 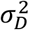 is the variance of the noise of the detector, and < *I* > is the average fluorescence intensity. For a photon-counting detector as used here, the brightness is ^64,95^:

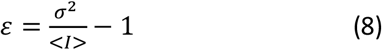

The brightness values, calculated for thousands of regions of interest, were potted as histograms.

## Acknowledgements

We thank Patrick Weber and Cédric Pissis the Crystallogenesis core facility at the Institut Pasteur for assistance with crystallization trials, and to the staff at the beamlines Proxima 1 and Proxima 2 at the French national synchrotron facility (SOLEIL, St Aubin, France) in particular to Leo Chavas and Bill Shepard for help with data collection and processing. We are grateful to Ignacio Fernandez and Jan Hellert for reading the manuscript and for their suggestions. This project has been supported via the recurrent funding from Institut Pasteur (MB, FR, PGC, RP, DB) and grants from NIH GM068619 and NSF MCB 1712740 (TL, KH).

## Contributions

Conceived and designed the experiments (MB, KH, FR), produced reagents (MB, DB, AH), collected the data (MB, PGC, TL), performed the data analyses (TL, PGC, RP, KH, MB), wrote the paper (MB, KH, TL, FR, PGC, RP, AH, FN).

