## Supplementary Information for "Human herpesvirus 8 molecular mimicry of ephrin ligands facilitates cell entry and triggers EphA2 signaling"

#### Table S1: Crystallographic statistics

| | **HHV-8 gH/gL – EphA2 LBD** |
|---|---|
| **Protein Data Bank code** | 7B7N |
| **Crystallization conditions** | |
| Protein conc. (mg/ml) | 5.1 |
| Crystallization buffer | 0.1M Na-malonate pH 5, 14.2% PEG 3350, 14 mM adenosine-5'-triphosphate disodium salt hydrate |
| Crystallization method | Sitting drop at 18°C |
| Cryo-protectant | 20% ethylene glycol |
| **Data Collection**[‡] | |
| Beamline | SOLEIL, Proxima 1 |
| Detector | Eiger X 16M |
| Space group | C 2 2 21 |
| Unit cell: a, b, c (Å) | 72.94, 129.00, 267.69 |
| α, β, γ (°) | 90, 90, 90 |
| Resolution (Å) | 46.06-2.69 (2.79-2.69) |
| Measured reflections | 479885 (44934) |
| Unique reflections | 35310 (3328) |
| Completeness (%) | 99.54 (96.24) |
| $CC_{1/2}$ (%)* | 99.7 (57.9) |
| Mean I/σ(I) | 9.13 (0.87) |
| Multiplicity | 13.6 (13.5) |
| B Wilson (Å$^2$) | 76.73 |
| Rsym | 0.2387 (2.117) |
| Rmeas | 0.2481 (2.199) |
| Rpim | 0.06717 (0.5884) |
| **Structure Determination** | |
| MR search models | EphA2 (PDB: 3HEI), EBV gH/gL (PDB: 3PHF) |
| N° of molecules in AU | 1 gH/gL-EphA2 LBD complex |
| **Refinement**[‡] | |

| | |
|---|---|
| Resolution cut-off (Å) | 46.06-2.69 (2.79-2.69) |
| Rwork (%) / Rfree (%) | 21.6 / 24.2 (21.1 / 25.4) |
| N° of Work / Free reflections | 33519 / 1762 (3161 / 165) |
| <B> atomic factors ($Å^2$) | 76.67 |
| N° of protein atoms | 7082 |
| N° of solvent atoms/ions | 54 / 98 |
| rmsd from ideal: | |
| Bond lengths (Å) | 0.003 |
| Bond angles (°) | 0.55 |
| Ramachandran[||] |
| Favored (%) | 95.47 |
| Allowed (%) | 4.08 |
| Outliers (%) | 0.45 |

[‡]Highest resolution shell is shown in parenthesis

*$CC_{1/2}$ is the correlation coefficient [1]

[||]Ramachandran values from MolProbity [2]

### Table S2: Interfaces between gH, gL and EphA2 LBD

The total surface area and the area at the interface, along with the number of atoms ($N_{at}$) and residues ($N_{res}$) are indicated for each pair of molecules. ΔG corresponds to the solvation free energy gain upon formation of the interface. Hydrogen bond distances cut-off of 3.5 Å, and 4.0 Å for salt bridges was applied; the number of hydrogen bonds and salt bridges are indicted with $N_{HB}$ and $N_{SB}$, respectively. Residues forming salt bridges are indicated with bold letters. The contacts made by residue E52$^{gH}$ are shown in blue. The interface analyses were done in PDBePISA [3].

#### A) gL interface with EphA2 LBD

| gL Surface (Å$^2$) | Interface area (Å$^2$) | $N_{at}$ | $N_{res}$ | ΔG (kcal/mol) | $N_{HB}$ | $N_{SB}$ |
|---|---|---|---|---|---|---|
| 7554 | 834 | 77 | 22 | -8.8 | 14 | 0 |

**Inter-chain contacts**

| ## | gL | Dist. [Å] | EphA2 |
|---|---|---|---|
| 1 | L:ALA 31[ N ] | 3.47 | E:ASP 61[ OD1] |
| 2 | L:SER 32[ N ] | 3.18 | E:ASP 61[ OD1] |
| 3 | L:SER 32[ OG ] | 2.95 | E:ASP 61[ OD1] |
| 4 | L:ARG 63[ NH2] | 3.71 | E:MET 55[ SD ] |
| 5 | L:THR 70[ N] | 3.00 | E:LEU 54[ O ] |
| 6 | L:GLU 72[ N] | 3.01 | E:GLN 56[ O ] |
| 7 | L:VAL 22[ O ] | 2.84 | E:ARG 103[ NH1] |
| 8 | L:VAL 22[ O] | 3.81 | E:CYS 188[ SG ] |
| 9 | L:GLN 30[ OE1] | 3.09 | E:ASN 60[ N ] |
| 10 | L:GLN 30[ OE1] | 3.13 | E:ASP 61[ N ] |
| 11 | L:ASP 68[ OD1] | 3.37 | E:TYR 48[ OH ] |
| 12 | L:THR 70[ O] | 3.01 | E:GLN 56[ N ] |
| 13 | L:GLU 72[ OE1] | 3.03 | E:ASN 57[ ND2] |
| 14 | L:ASN 128[ OD1] | 3.83 | E:ASN 60[ N ] |

#### B) gH interface with EphA2 LBD

| gH Surface ($Å^2$) | Interface area ($Å^2$) | $N_{at}$ | $N_{res}$ | ΔG (kcal/mol) | $N_{HB}$ | $N_{SB}$ |
|---|---|---|---|---|---|---|
| 29843 | 121.4 | 15 | 4 | -0.4 | 2 | 1 |

**Inter-chain contacts**

| ## | gH | Dist. [Å] | EphA2 |
|---|---|---|---|
| 1 | H:GLU 52[ O ] | 2.91 | E:ARG 103[ NH2] |
| **2** | **H:GLU 52[ OE1]** | **2.94** | **E:ARG 103[ NH1]** |

#### C) gL interface with gH

| gL Surface ($Å^2$) | Interface area ($Å^2$) | $N_{at}$ | $N_{res}$ | ΔG (kcal/mol) | $N_{HB}$ | $N_{SB}$ |
|---|---|---|---|---|---|---|
| 7554 | 2385 | 264 | 63 | -36.3 | 24 | 3 |

**Inter-chain contacts**

| ## | gL | Dist. [Å] | gH |
|---|---|---|---|
| 1 | L:ILE 46[ N ] | 2.90 | H:SER 48[ O ] |
| 2 | L:PHE 48[ N ] | 2.91 | H:GLU 50[ O ] |
| 3 | L:VAL 22[ N ] | 3.01 | H:GLU 52[ OE1] |
| 4 | L:HIS 47[ ND1] | 2.56 | H:GLU 52[ OE2] |
| 5 | L:ASN 79[ ND2] | 3.49 | H:ALA 81[ O ] |
| 6 | L:ASN 79[ ND2] | 2.69 | H:GLU 82[ O ] |
| 7 | L:ASN 76[ N ] | 2.48 | H:VAL 83[ O ] |
| 8 | L:ASN 76[ ND2] | 3.00 | H:GLU 85[ O ] |
| 9 | L:ASN 79[ ND2] | 3.18 | H:GLU 85[ OE2] |
| 10 | L:ASN 76[ ND2] | 3.00 | H:THR 90[ OG1] |
| 11 | L:SER 82[ OG ] | 2.66 | H:TYR 172[ OH ] |
| 12 | L:ARG 90[ NH1] | 3.03 | H:PRO 173[ O ] |
| 13 | L:ARG 89[ NH2] | 2.89 | H:ASP 227[ O ] |
| 14 | L:ARG 89[ NH2] | 2.68 | H:LEU 229[ O ] |
| 15 | L:ARG 89[ NH1] | 3.54 | H:SER 231[ OG ] |
| 16 | L:PRO 38[ O ] | 2.99 | H:TRP 78[ NE1] |
| 17 | L:PHE 41[ O ] | 2.76 | H:ARG 44[ NH2] |
| 18 | L:VAL 43[ O ] | 2.88 | H:ARG 44[ NH1] |
| 19 | L:HIS 44[ O ] | 3.55 | H:SER 48[ OG ] |
| 20 | L:HIS 44[ O ] | 3.68 | H:SER 48[ N ] |
| 21 | L:ILE 46[ O ] | 2.65 | H:GLU 50[ N ] |
| 22 | L:PHE 48[ O ] | 2.95 | H:GLU 52[ N ] |

| | | | |
|---|---|---|---|
| 23 | L:ASN 79[ O ] | 3.36 | H:TYR 172[ OH ] |
| 24 | L:ASP 123[ OD1] | 3.67 | H:ARG 88[ NH2] |
| **25** | **L:HIS 47[ NE2]** | **3.67** | **H:GLU 50[ OE1]** |
| **26** | **L:HIS 47[ ND1]** | **2.56** | **H:GLU 52[ OE2]** |
| **27** | **L:ASP 123[ OD1]** | **3.67** | **H:ARG 88[ NH2]** |

#### Table S3: Properties of the dimerization interface (DIN) in EphA2 in unliganded form and bound to ligands

| | Surface (Å²) | Interface area (Å²) | $N_{at}$ | $N_{res}$ | $\Delta G$ (kcal/mol) | $N_{HB}$ | $N_{SB}$ |
|---|---|---|---|---|---|---|---|
| **EphA2 ecto** PDB: 3FL7 | 25735 | 424 | 51 | 16 | -3.6 | 8 | 0 |
| **EphA2 ecto + ephrin-A5** PDB: 2X11 | 26737 | 532 | 61 | 17 | -5.3 | 6 | 0 |
| **EphA2 LBD-CRD + ephrin-A1** PDB: 3MBW | 15500 | 537 | 61 | 19 | -4.2 | 10 | 0 |
| **EphA2 LBD + ephrin A1** PDB: 3CZU | 9301 | 634 | 70 | 17 | -6.9 | 7 | 0 |
| **EphA2 LBD + HHV8 gH/gL** PDB: 7B7N | 8381 | 693 | 83 | 21 | -4.9 | 16 | 0 |

#### Figure S1: Secondary structure topology diagram of EphA2 LBD, gL and ephrin-A1

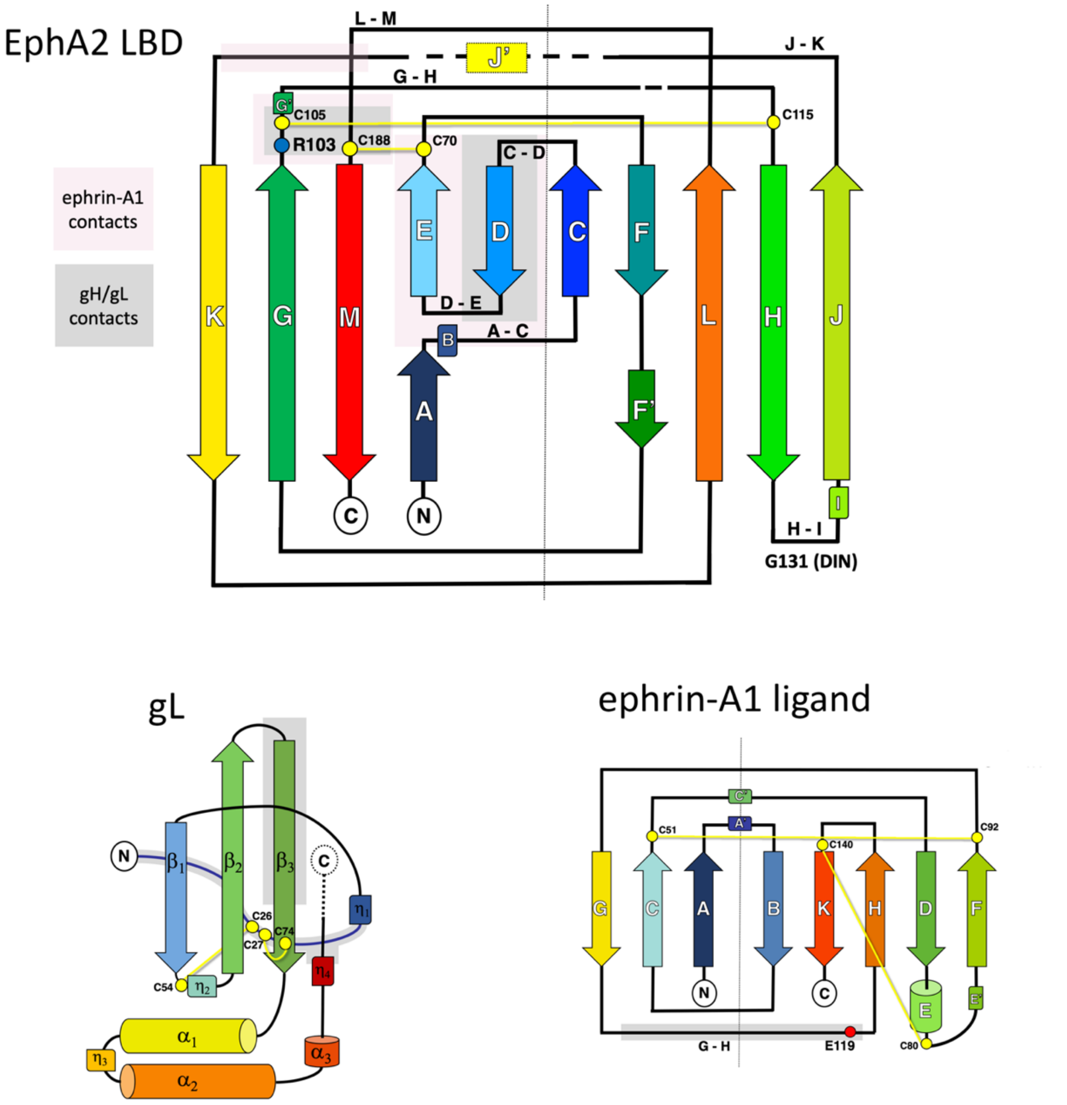

**Figure S1 legend**: EphA2 LBD - grey and pink shaded areas indicate the structural elements involved in interactions with gH/gL and ephrin-A1, respectively. In gL and ephrin-A1 grey shaded areas highlight the structural elements involved in interactions with EphA2 LBD. Secondary structure elements are represented by arrows (β-strands), rectangles (α-helices) and rounded rectangles (η helices (B, I, J')). The horizontal dashed lines indicate residues missing from the structure. The vertical dotted line designates the two 5-stranded β-sheets adopting a jelly-roll fold in EphA2 LBD and a 3- and 5-stranded sheets forming a β-sandwich in ephrin-A1. The conserved residues $R103^{EphA2}$ and $E119^{ephrin\text{-}A1}$, which are important for high affinity interaction are represented as a red and blue circle, respectively.

### Figure S2: Oligomeric assemblies formed by EphA2 ectodomains

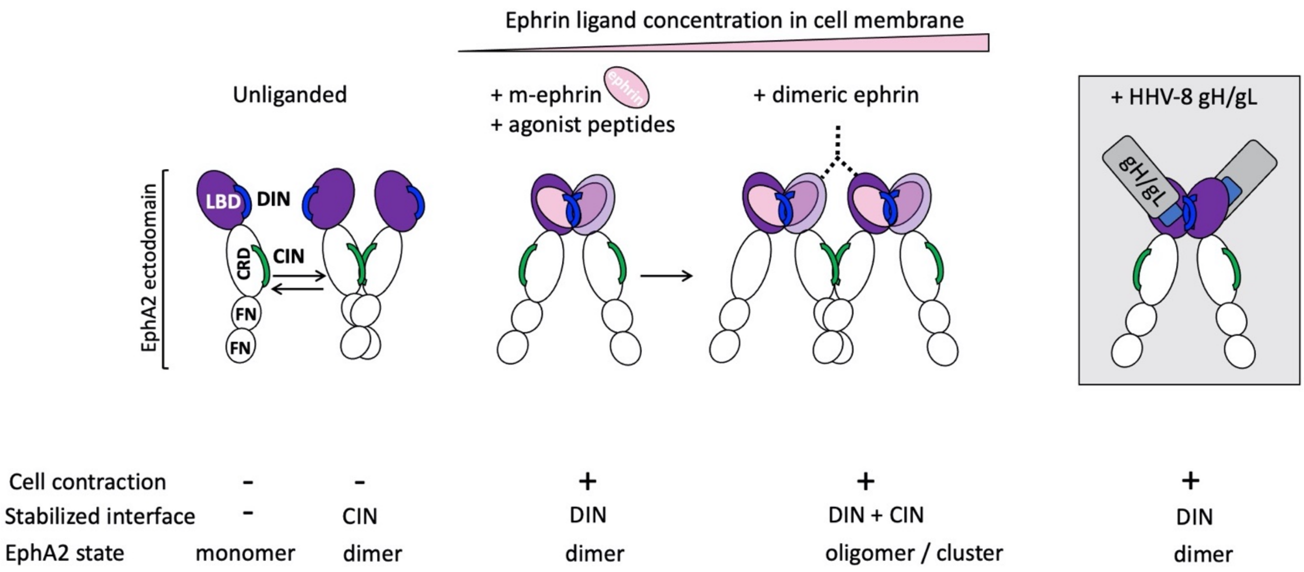

**Figure S2 legend:** the EphA2 ectodomain is represented with the LBD colored in purple and the cysteine rich domain (CRD) and 2 fibronectin domains (FN) in white. The dimerization interface (DIN) in LBD and clustering interface (CIN) in CRD are indicated as blue and green, respectively. Ephrin ligand is depicted with a pink oval, and gH/gL as grey and blue rectangles.

### Figure S3: EphA2 LBD binds to HHV-8 gH/gL in 1:1 stoichiometry

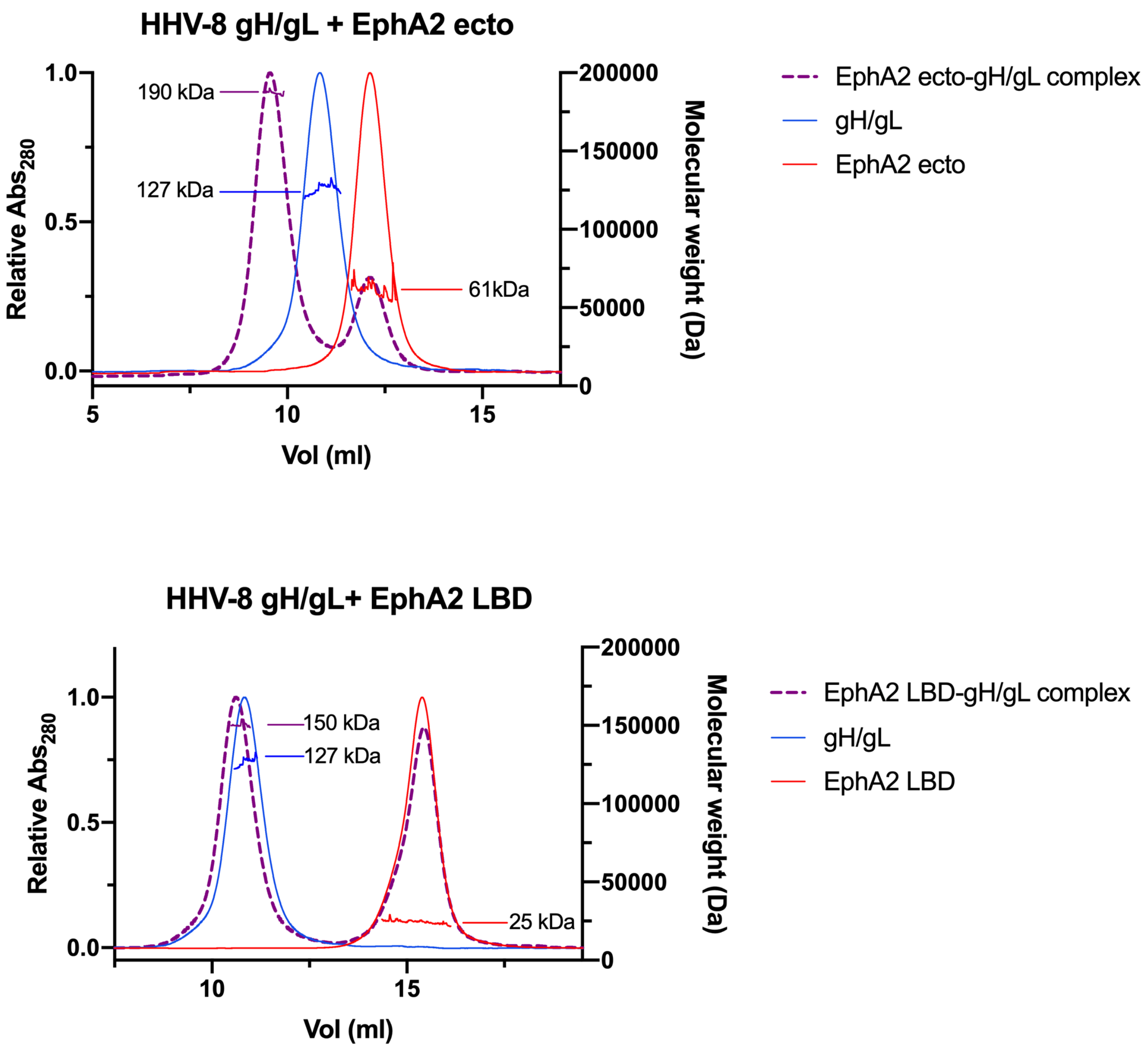

**Figure S3 legend**: MALS measurements of gH/gL (blue curve), EphA2 ectodomain or LBD (red curve), and gH/gL-EphA2 complexes (purple, dashed curve). Molecular weights are indicated on the chromatograms, demonstrating that the tertiary complex is composed of 1 molecule of gH/gL bound to 1 molecule of EphA2 ectodomain or LBD.

#### Figure S4: Structural comparison of gH/gL from gamma- (HHV-8, EBV), beta- (CMV) and alpha-herpesviruses (HSV2 and VZV)

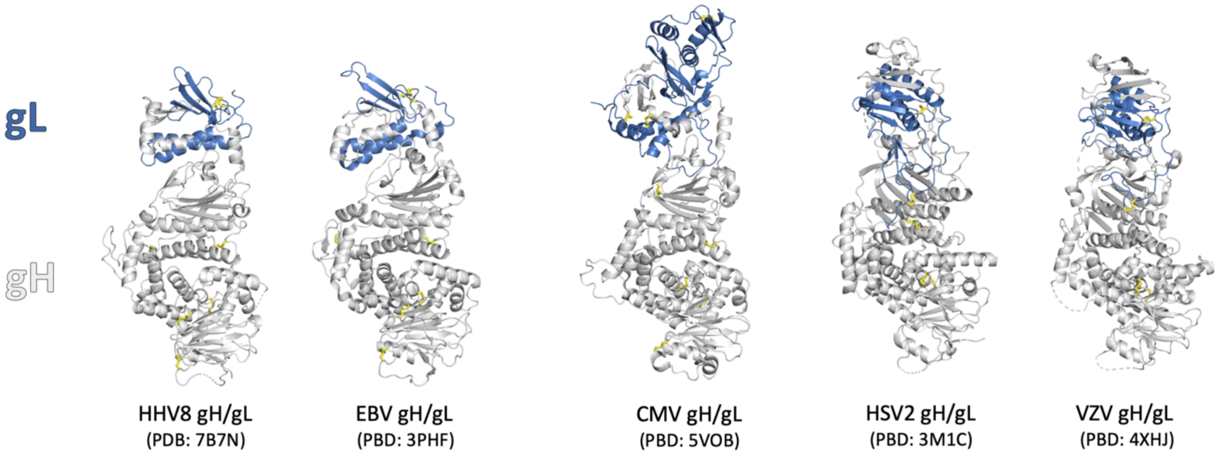

| | | EBV | | | | CMV | | | | HSV-2 | | | | VZV | | | |
|---|---|---|---|---|---|---|---|---|---|---|---|---|---|---|---|---|---|
| HHV-8 | | Z | rmsd | lali/nres | %id | Z | rmsd | lali/nres | %id | Z | rmsd | lali/nres | %id | Z | rmsd | lali/nres | %id |
| gH | DI | 3.1 | 2.7 | 43/46 | 26 | - | - | - | - | - | - | - | - | - | - | - | - |
| | DII | 28.1 | 2.7 | 262/279 | 21 | 19.3 | 3.1 | 230/282 | 17 | 10.2 | 3.3 | 148/210 | 9 | 10.1 | 3.3 | 163/206 | 7 |
| | DIII | 18.4 | 2.3 | 171/185 | 25 | 14.6 | 2.8 | 154/168 | 14 | 13.6 | 2.8 | 152/179 | 18 | 12.0 | 3.1 | 150/184 | 18 |
| | DIV | 17.3 | 2.4 | 132/143 | 39 | 15.2 | 2.3 | 129/145 | 22 | 15.1 | 2.3 | 125/148 | 20 | 14.5 | 2.5 | 131/150 | 19 |
| gL | | 12.3 | 1.7 | 97/108 | 24 | 2.6 | 3.8 | 75/237 | 15 | 6.6 | 3.2 | 94/146 | 11 | 6.3 | 3.3 | 93/132 | 13 |

**Figure S4 legend**: The structural alignments were performed using Dali Pairwise Structure Comparison server [4], with the PDB coordinate files noted on the figure. The domain boundaries were defined based on the sequence alignments; because of the variability in the length of the gH DI and poor or no conservation at the amino acid level, the hinge/linker helix was used as a demarcation point for the boundary between DI and DII.

Z-scores are calculated as reported in [4] and indicate structural similarity. 'Rmsd' (root-mean-square deviation) is the average distance deviation between the aligned C$\alpha$ atoms in Å. 'lali' refers to the number of aligned i.e. structurally equivalent residues, 'nres' is the total number of residues in the target protein.

### Figure S5: Analyses of gH/gL and ephrin-A1 interfaces with EphA2 LBD

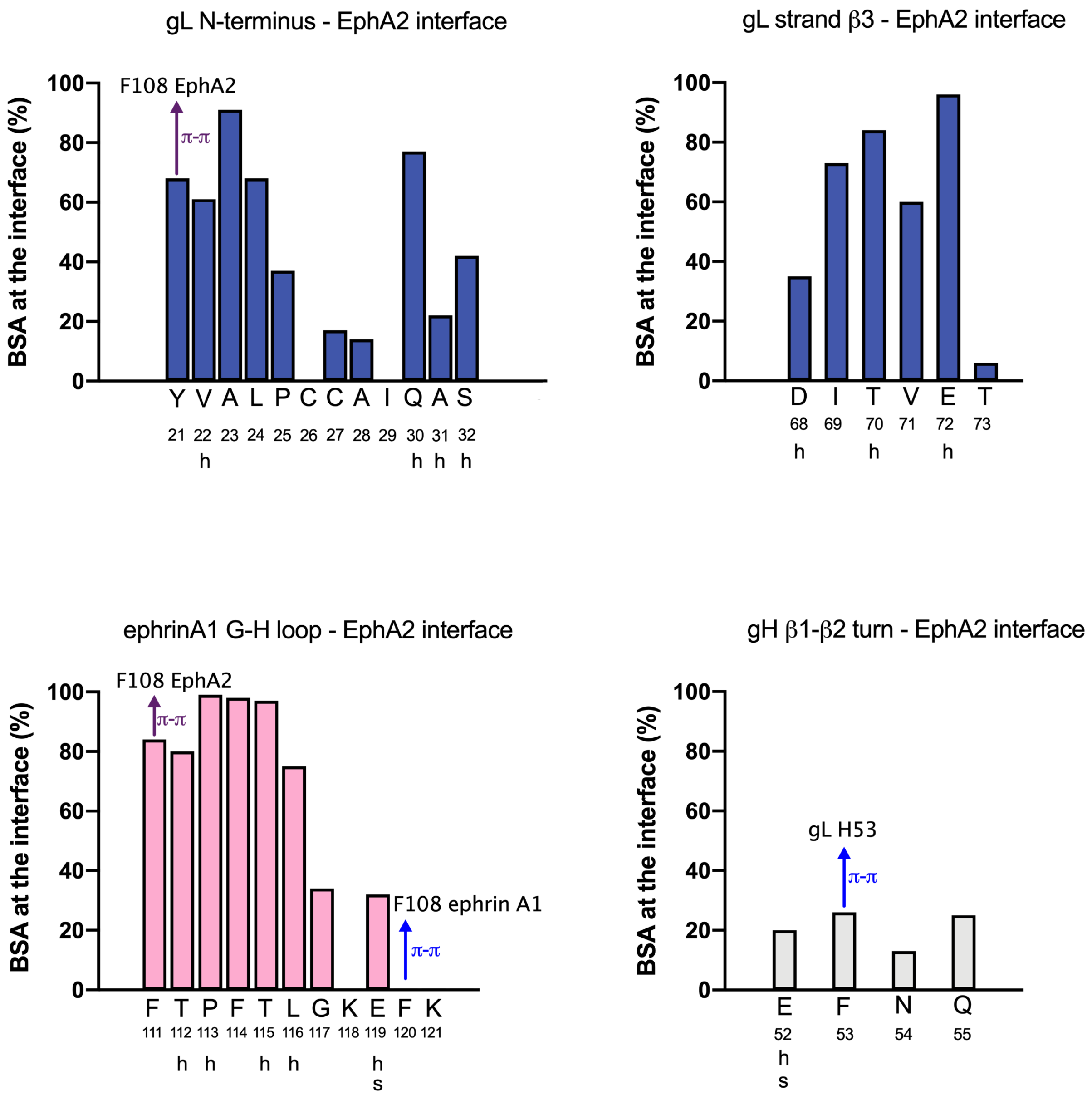

**Figure S5 legend**: The buried surface area (BSA) is presented as % of the total residue surface, and is plotted for each residue, indicated by a letter and number on the x-axis, for each given interface. The residues participating in hydrogen and salt bridge bonds are marked with 'h' and 's', respectively. The residues involved in pi-pi interactions are labeled with blue arrows, and their contact residues are indicated.

#### Figure S6: Illustration of the BLI setup

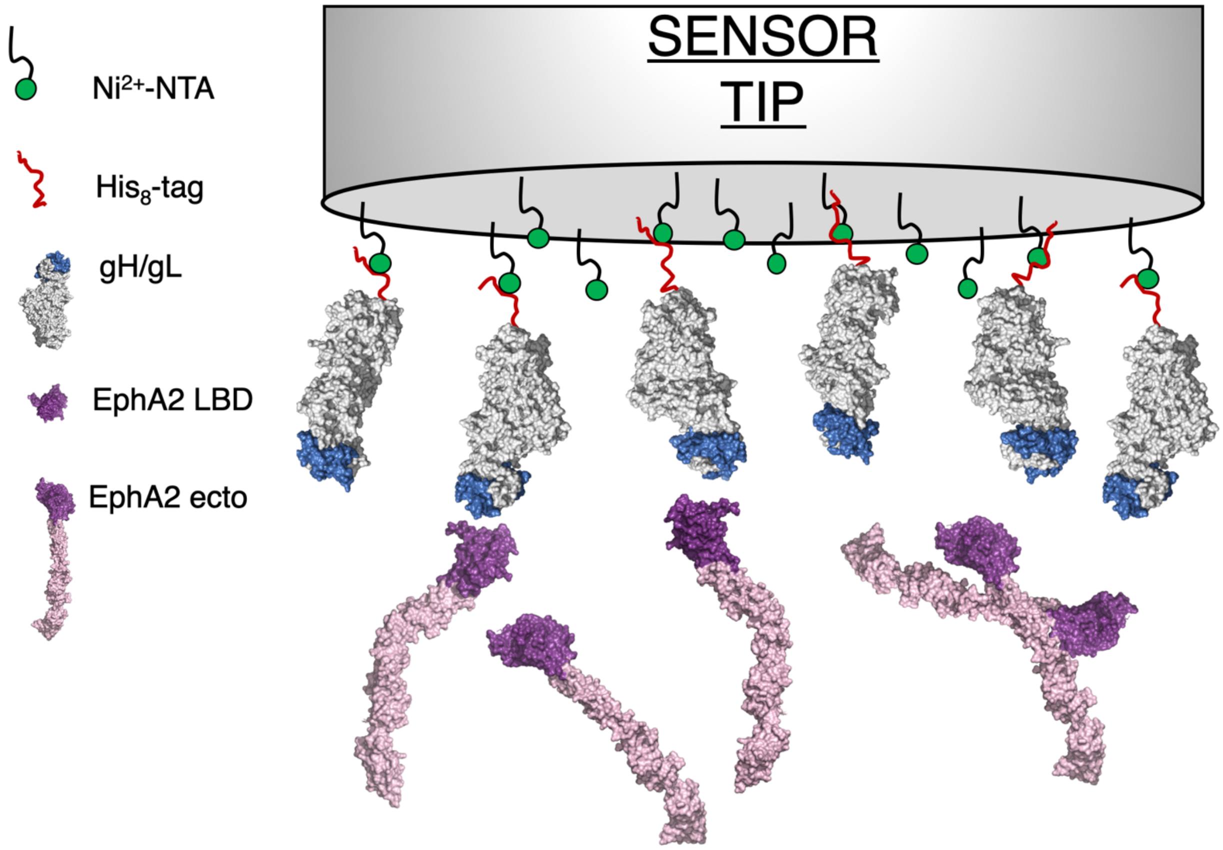

**Figure S6 legend**: HHV-8 gH/gL is loaded onto the NTA-Ni$^{2+}$ sensors via a histidine tag attached to the gH C-terminus located at the opposite side from the gL, and the EphA2 binding site.

#### Figure S7: Binding of WT gH/gL and EphA2 at low pH and in inverted system

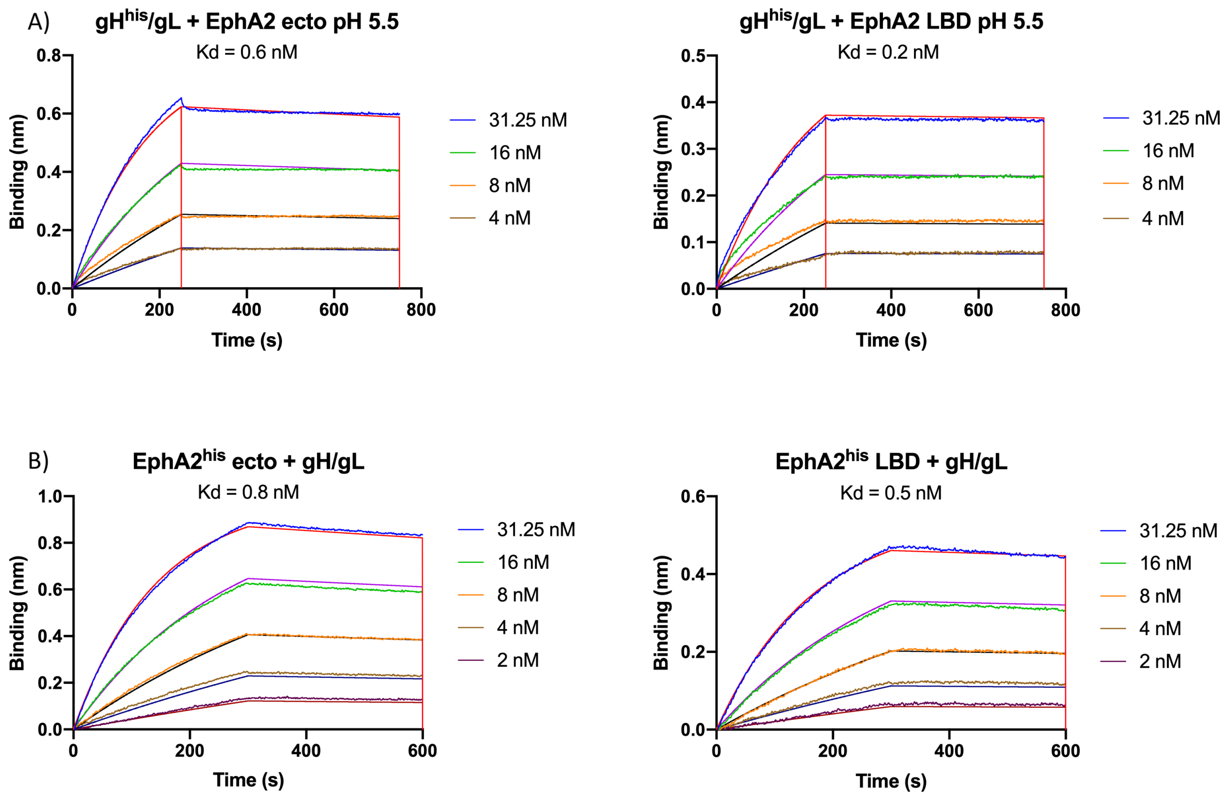

**Figure S7 legend**: BLI sensorgrams obtained for A) interactions between immobilized gH/gL and EphA2 ectodomain / LBD at pH 5.5, and B) interactions between immobilized EphA2 ectodomain / LBD and gH/gL at pH 7.5.

### Figure S8: FSI-FRET data: Proximity-corrected FRET efficiencies, donor concentrations, and acceptor concentrations.

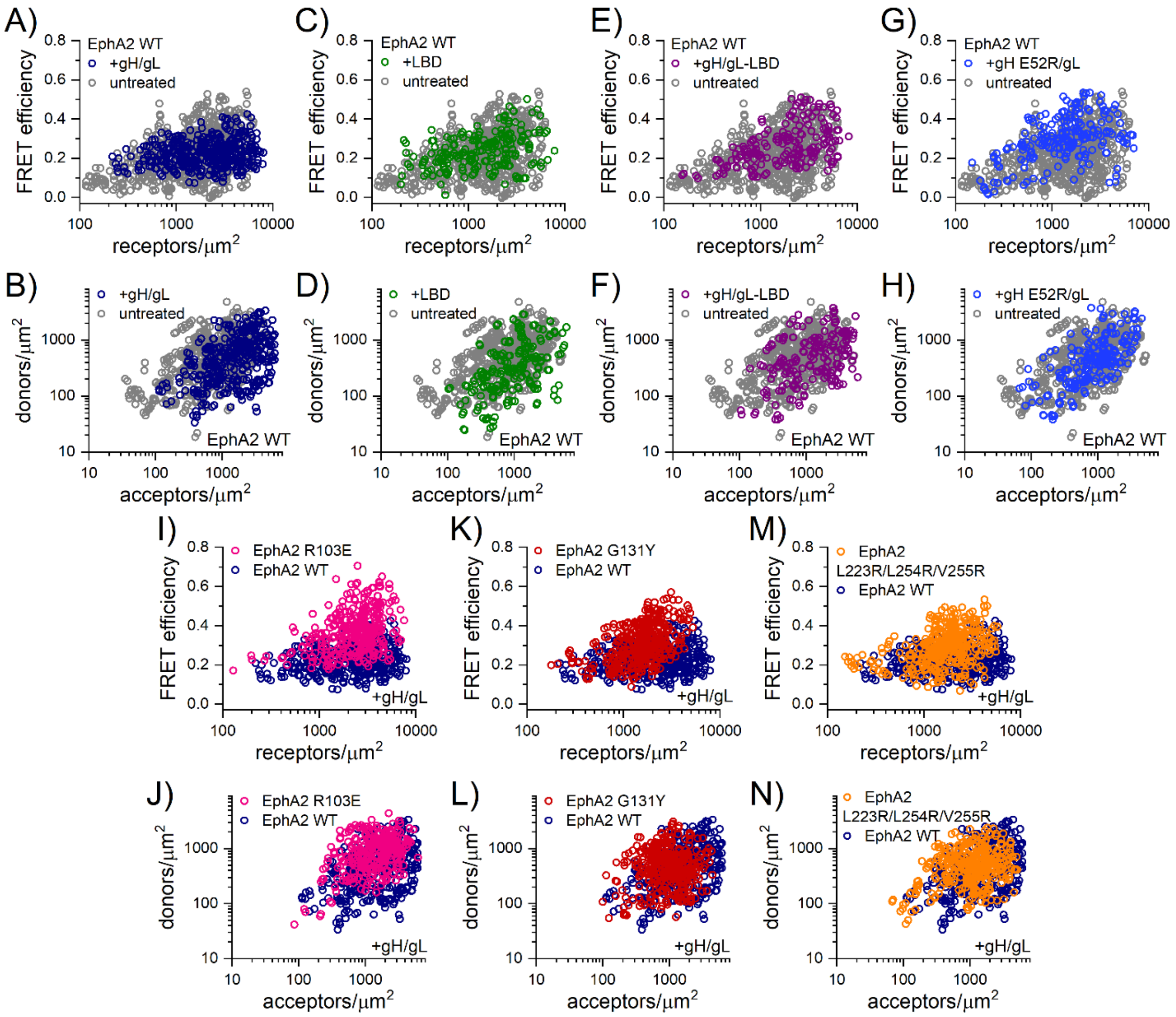

**Figure S8 legend**: The FSI-FRET method determines the FRET efficiencies, the concentration of donor-tagged EphA2 (EphA2-mTURQ), and the concentration of acceptor-tagged EphA2 (EphA2-eYFP) at the plasma membrane of live HEK293T cells. The FRET efficiencies were corrected for the non-specific 'proximity FRET' contribution and are plotted as a function of the measured receptor concentration (EphA2-mTURQ+EphA2-eYFP concentrations). The proximity-corrected FRET efficiencies and the donor and acceptor concentrations were measured for the following conditions: (**A-B**) EphA2 WT +gH/gL, (**C-D**) EphA2 WT +LBD, (**E-F**) EphA2 WT +gH/gL-LBD, (**G-H**) EphA2 WT +gH E52R/gL, (**I-J**) EphA2 R103E +gH/gL, (**K-L**) EphA2 G131Y +gH/gL, (**M-N**) EphA2 L223R/L254R/V255R +gH/gL. The data in (**A-H**) are compared to EphA2 WT data in the absence of ligand, which was previously reported [5]. The data in (**I-N**) are compared to EphA2 WT in the presence of gH/gL (from (**A-B**)).

#### Figure S9: EphA2 assemblies and contacts observed in the crystal

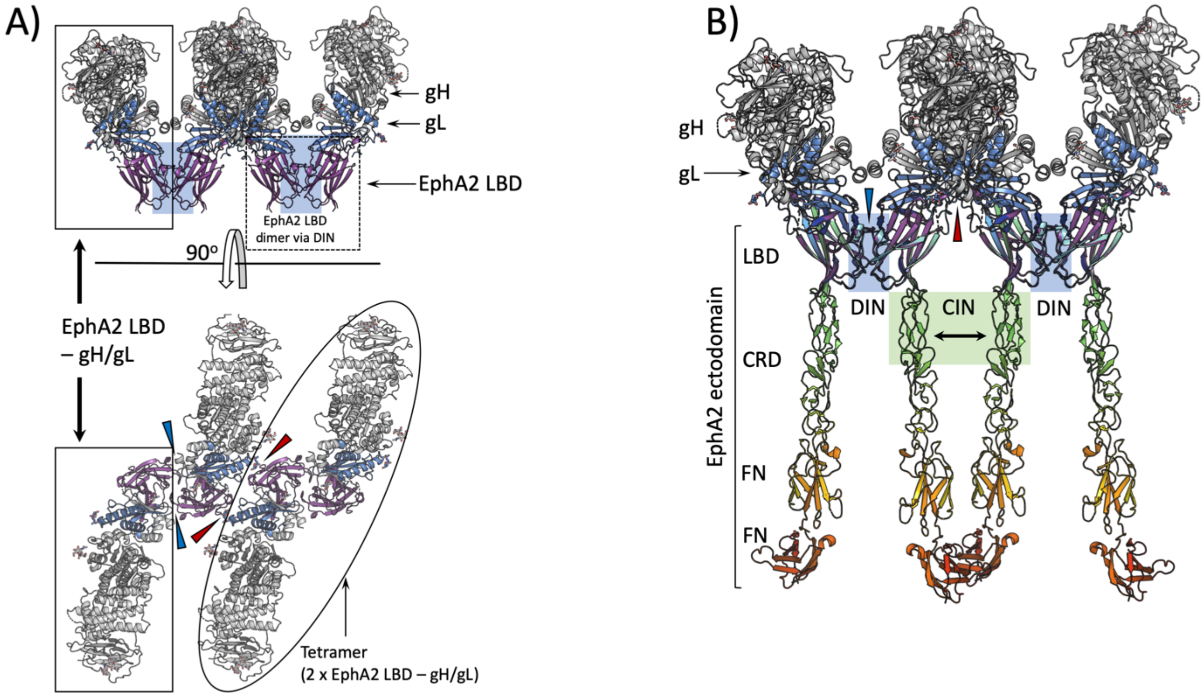

**Figure S9 legend: A)** Crystal packing of EphA2 LBD – gH/gL complexes. The solid line rectangles indicate a single tertiary complex (gH/gL-EphA2 LBD). The EphA2 LBD dimer formed via DIN (blue shaded box) is enclosed with a dashed-line rectangle. Bottom panel shows the top view; a tetramer formed of 2 gH/gL and 2 EphA2 LBD molecules is indicated with an oval shape. The blue triangles indicate absence of contacts between gH/gL of one tertiary complex with the LBD from the adjacent complex molecule within the same tetramer. Red triangles point to the sites of contacts of gL with LBD from another tetrameric assembly. The latter contacts are formed between the NAG moiety at the N118 of gL and res C115 of EphA2 LBD (1 HB). **B)** The structure of the EphA2 ectodomain (PDB 2X10, rainbow colors) was superimposed onto the EphA2 LBD bound to gH/gL to indicate the putative location of the EphA2 domains downstream from LBD in this kind of arrangement. The DIN and CIN are marked with the blue and green boxes, respectively. The distance between CIN, imposed by gH/gL, is too large for contacts to be established and drive aggregation of EphA2 dimers into larger oligomers.

### Figure S10: Alignment of gL sequences from gammaherpesviruses

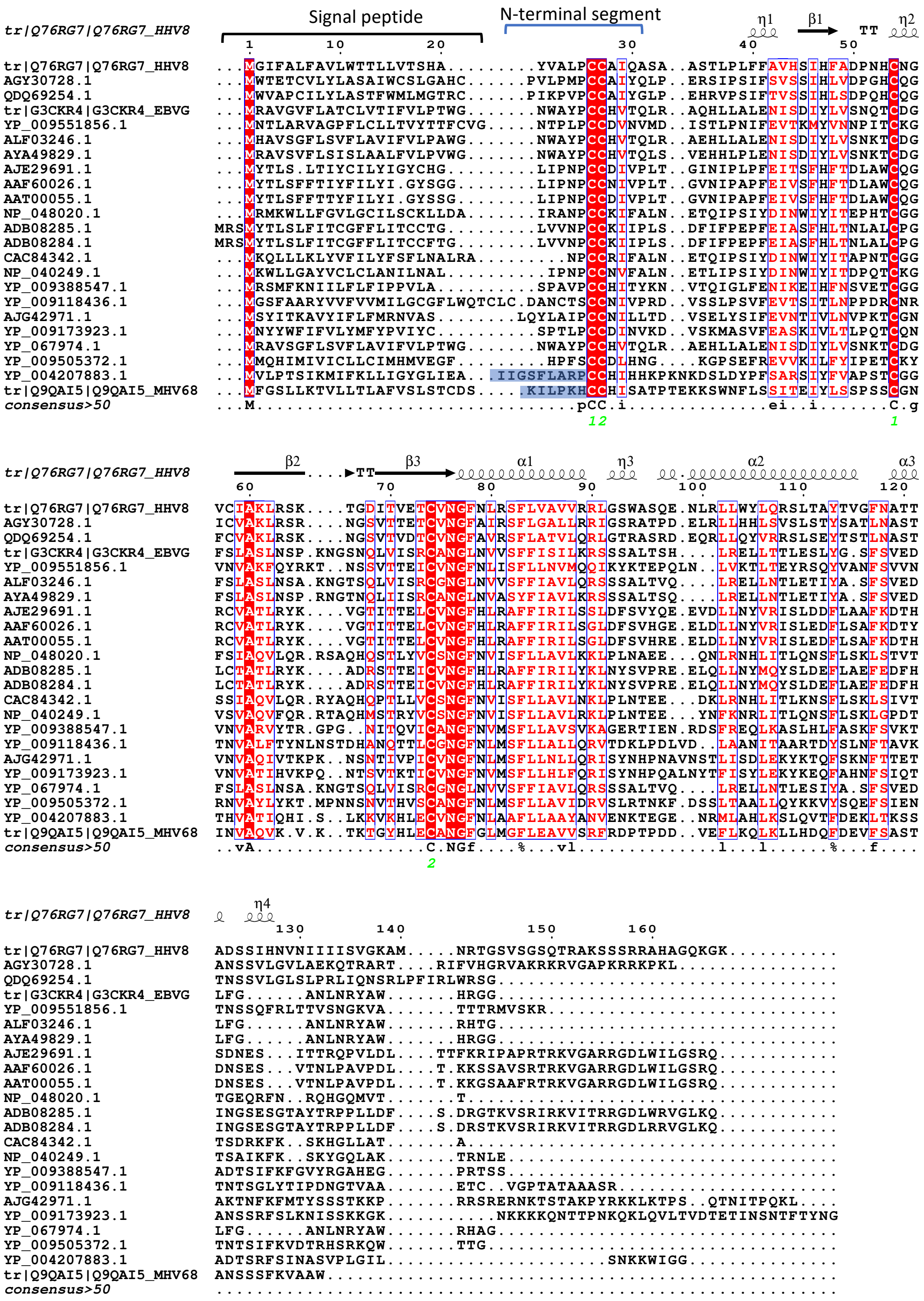

**Figure S10 legend:** The HHV-8 gL sequence is placed on the top. The N-termini in two rodent gLs (Cricetid gammaherpesvirus 2, accession number YP_004207883.1, and Murine gammaherpesvirus 68, accession number Q9QAI5_MHV68) contain positively charged residues and are shaded in blue on the bottom of the alignment. Secondary structure elements are indicated above the sequences, and the disulfide bridges (green letters) and consensus sequence below. The alignment was generated by Clustal Omega[6] and plotted by ESPript [7].
